# Transcription-dependent domain-scale 3D genome organization in dinoflagellates

**DOI:** 10.1101/2020.07.01.181685

**Authors:** Georgi K. Marinov, Alexandro E. Trevino, Tingting Xiang, Anshul Kundaje, Arthur R. Grossman, William J. Greenleaf

## Abstract

Dinoflagellate chromosomes represent a unique evolutionary experiment, as they exist in a permanently condensed, liquid crystalline state, are not packaged by histones, and contain genes organized into polycistronic arrays, with minimal transcriptional regulation. We analyze the 3D genome of *Breviolum minutum*, and find large topological domains without chromatin loops, demarcated by convergent gene array boundaries (“dinoTADs). Transcriptional inhibition degrades dinoTADs, implicating transcription-induced supercoiling as the primary topological force in dinoflagellates.

---

The three-dimensional (3D) genome architecture of cells has functional consequences for gene regulation, organismal development, replication, and mutational processes. Mechanisms known to drive genome folding in eukaryotes include constraints on cohesin-mediated loop extrusion – imposed by CTCF in vertebrates – that generate topologically associating domains (TADs), and self-associations between similar chromatin states that form compartments ^1^. However, the extent to which genome function itself may influence genome folding, for example through transcriptional activity, is poorly understood. There has also been little exploration of 3D organization across eukaryotes, even though major deviations from conventional norms are known to exist, presenting natural experiments that may reveal deeper underlying organizational principles masked in other lineages.

Dinoflagellates are the most radical such departure. They are a diverse, widespread clade playing major roles in aquatic ecosystems, for example, as symbioints of corals, providing the metabolic basis for reef ecosystems. Dinoflagellates possess numerous highly divergent molecular features ^2^, including, uniquely among eukaryotes, the loss of nucleosomal packaging of chromatin. Histones are extremely conserved across eukaryotes, were present in their current form already in the Last Eukaryotic Common Ancestor ^3^, and they and their posttranslational modifications are pivotal to all biochemical processes involving chromatin.

Dinoflagellates are the sole known exception. Their chromosomes exist in a liquid crystalline state, are permanently condensed throughout the cell cycle, and, although highly divergent histone genes are retained in their genomes ^4^, a combination of virus-derived nucleoproteins and bacterial-derived histone-like proteins have taken over as main packaging components ^5^. Dinoflagellate genomes are often huge (up to ≥ 100 Gbp), genes are organized into polycistronic gene arrays, individual mRNAs are generated through *trans*-splicing, and transcriptional regulation is largely absent. These fascinating features simultaneously pose intriguing questions regarding the adaptation of transcriptional and regulatory mechanisms to the absence of nucleosomes, and provide a unique opportunity to explore the biophysical forces underlying genomic organization in the context of a large eukaryotic genome nearly devoid of nucleosomes.

To explore these questions, we performed Hi-C on the coral symbiont *Breviolum minutum*. We generated multiple libraries under standard growth conditions and for cells grown at elevated temperature, obtaining ∼ 150–220 million Hi-C contacts for each (Supplementary Table 1). We pooled these libraries to generate a chromosome-level scaffolding of the previously fragmented *B. minutum* assembly ^6^. We identified 91 major pseudochromosomes (≥ 500 kbp), encompassing ∼94% of the total sequence (Fig. 1A-B; Supplementary Fig. 1A), the longest being ∼11 Mbp in size, with a median length of 6.7 Mbp (Supplementary Fig. 1A). At 1-Mbp resolution, they exhibit a bipartite (occasionally tripartite) structure (Supplementary Fig. 2).

**Figure 1:**
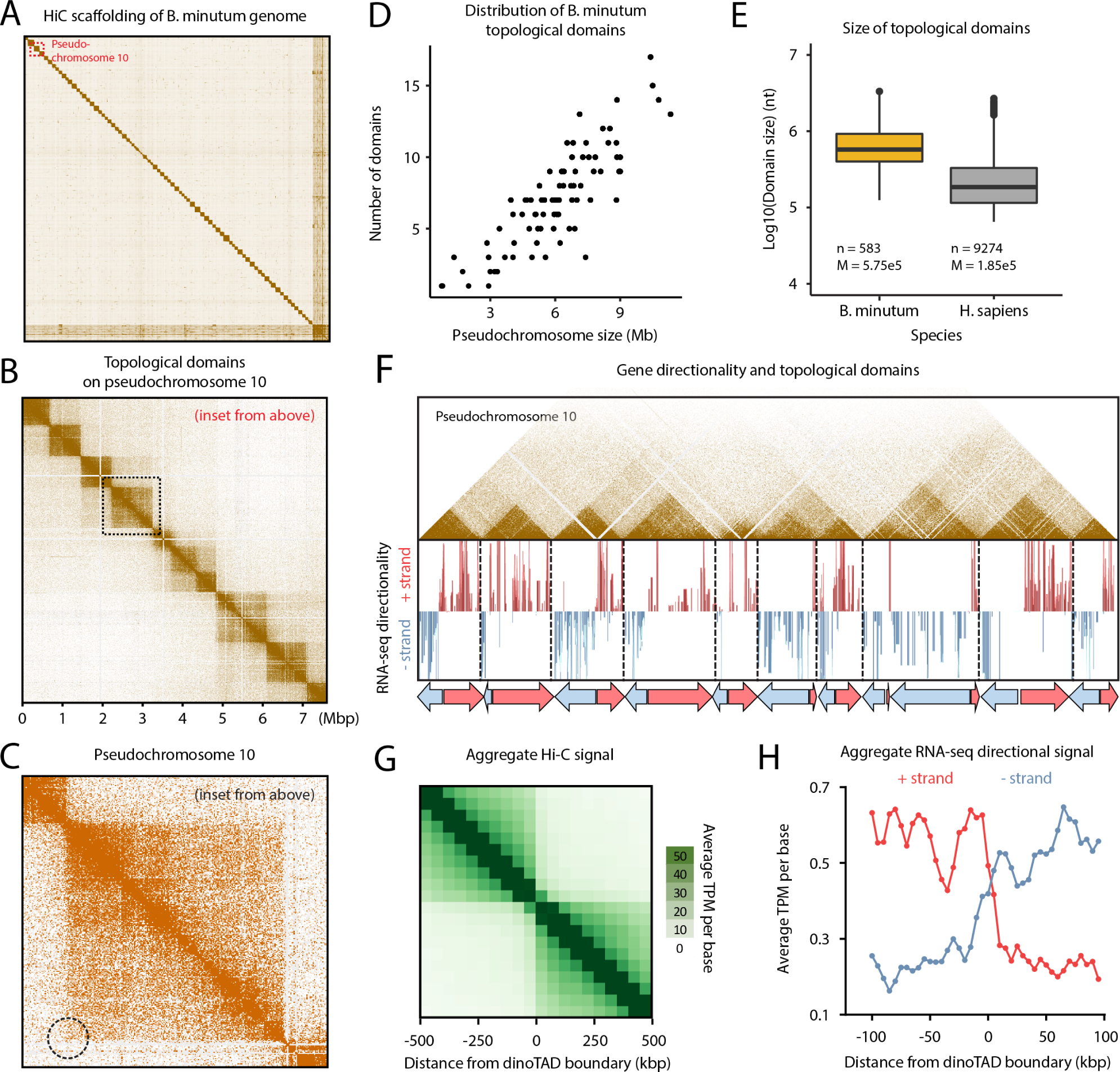
*B. minutum* genome is physically partitioned into dinoTADs defined by polycistronic gene arrays. (A) Hi-C scaffolding of the *B. minutum* draft genome assembly. (B) Inset from (A). KR-normalized 5-kb resolution Hi-C map for pseudochromosome 10. (C) Inset from (B). Hi-C loops and stripes are not observed in dinoTADs. (D) Scales of chromosome size with dinoTAD number. (E) Comparison of human and *B. minutum* topological domain sizes. (F) Hi-C map for pseudochromosome 10 together with forward- and reverse-strand transcript levels and gene arrays. (G) Average Hi-C contacts across dinoTAD boundaries. (H) Average forward- and reverse-strand RNA-seq levels across dinoTAD boundaries.

High-resolution maps revealed very strong topological domains,≤ 200– ≥ 2 Mbp in size (Fig. 1B-E; Supplementary Fig. 3–12). In mammals, TAD boundaries are demarcated by CTCF sites blocking loop extrusion, reflected in Hi-C maps by chromatin loops and “stripes”. We observe no loop or stripe features in *B. minutum* (Fig. 1C), suggesting a different mechanism for the formation of dinoflagellate TADs, which we term “dinoTADs”. DinoTAD number correlates with chromosome size (Fig. 1D), and they are considerably larger than mammalian TADs (Fig. 1E).

We next compared Hi-C maps to available annotation features. Remarkably, we found that each dinoTAD corresponds to a pair of divergent gene arrays (Fig. 1F), and dinoTAD boundaries coincide with convergence between gene arrays (Fig. 1G-H).

The correspondence between dinoTADs and gene arrays suggested a role for transcription in their formation. Although TADs form independently of transcription in metazoan cells, transcription-induced self-interacting domains have been previously demonstrated in bacteria ^8^, and similar mechanisms have been proposed to explain some topological features in fission yeast ^9^. This model makes a clear prediction – inhibition of transcription should result in dinoTADs decompaction.

To test this relationship, we first compared Hi-C maps for cells grown at 34 °C versus 27 °C, as heat stress could result in general transcription reduction ^10^. We observed mild decompaction of dinoTADs at 34 °C, though domains remained intact (Supplementary Fig. 18–20).

We next carried out chemical transcription inhibition experiments. Since transcription inhibition conditions for *B. minutum* are not well established, we chose two inhibitors – triptolide and *α*-amanitin – with distinct mechanisms of action, and assayed multiple time points and doses (Fig. 2A-B). Amanitin directly inhibits RNA Polymerase II and is slow acting, while triptolide quickly blocks initiation by targeting the TFIIH XPB subunit ^11^. However, the *B. minutum* XPB homolog is highly divergent^6^, thus a moderate inhibition effect is not unexpected. Indeed, while we observed clear dose-dependent blurring of dinoTAD boundaries after triptolide treatment, broad dinoTAD-like structures remained (Fig. 2E-F; Supplementary Figures 25–28).

**Figure 2:**
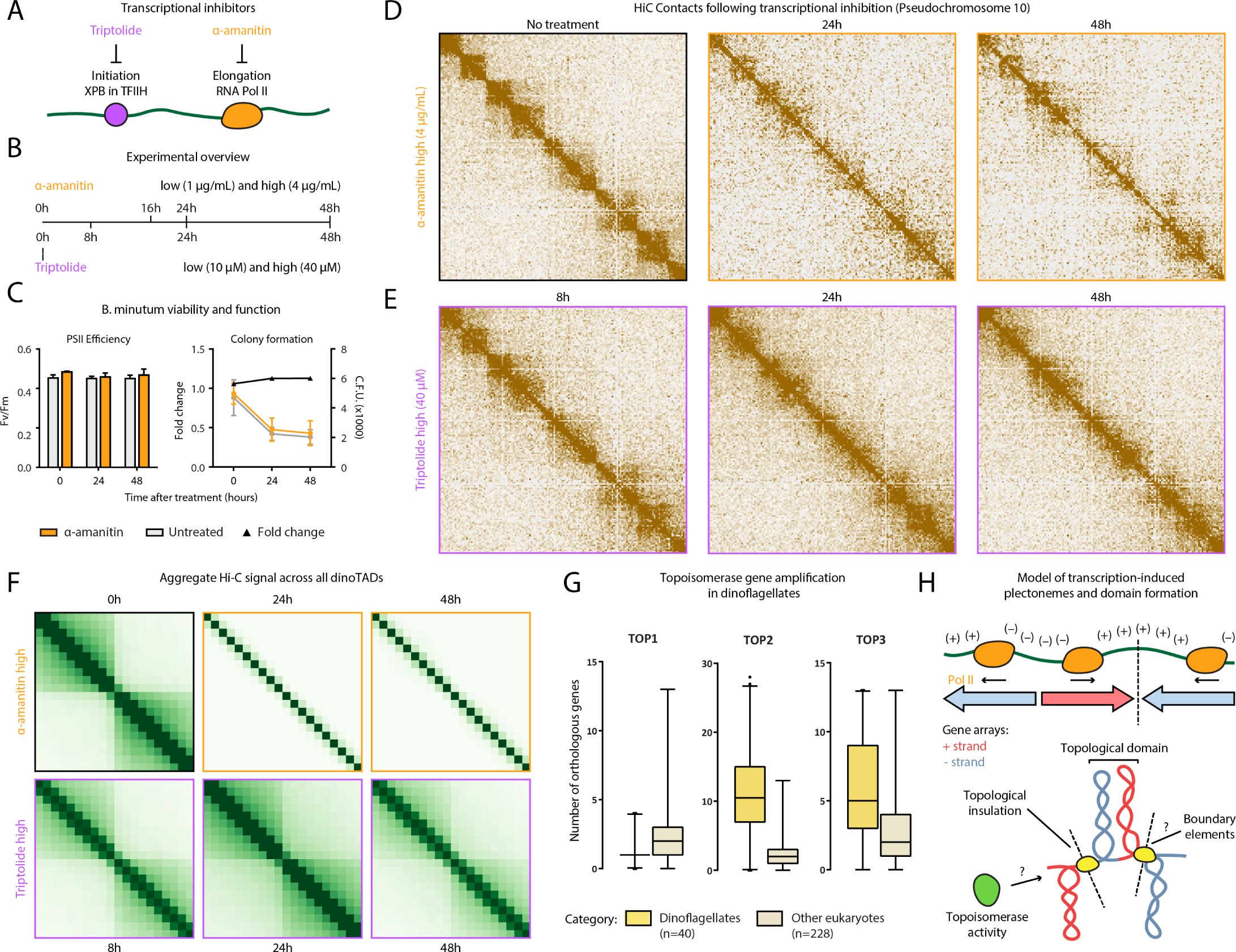
Decompaction of dinoTADs upon application of transcriptional inhibitors and the transcription-induced supercoiling model for their formation. Shown is pseudochromosome 10. (A-B) Outline of transcription inhibition time course experiments. (C) Comparison of cell function, measured by PSII photosythetic efficiency, and cell viability, measured by colony formation (right), between *α*-amanitin-treated and untreated cells. (D) KR-normalized Hi-C maps (50-kb resolution) show marked loss of dinoTADs after *α*-amanitin treatment. (E) Hi-C maps show reduction of insulation at dinoTAD boundaries after triptolide treatment. (F) Metaplots of Hi-C signal around domain boundaries (50-kb resolution). (G) Amplification of *TOP2* and *TOP3* topoisomerases in dinoflagellates (based on MMETSP ^7^ transcriptome assemblies). (H) Transcription-induced supercoiling as driver of dinoflagellate chromatin folding. Transcribing polymerases introduce negative/positive DNA supercoiling behind/ahead of the transcription machinery. Interactions within supercoiled domains could explain the physical association of divergently-oriented arrays. Topological insulation could be driven by supercoiling-related effects, or by specific boundary elements.

Strikingly, *α*-amanitin treatment resulted in a dosedependent, progressive, near-complete dinoTAD decompaction (Fig. 2D,F; Supplementary Fig. 21–24). These effects were observed in both technical and biological replicates (Supplementary Fig. 21–24).

Of note, even at high doses, *α*-amanitin treatment did not detectably affect photosynthetic efficiency or cell viability relative to controls (Fig. 2C), excluding cell death as a confounding factor.

These experiments support a transcription-induced supercoiling model for dinoTAD formation. Torque generated by active polymerases produces positive/negative supercoiling ahead of/behind the transcription bubble. This can alter the twist of the double helix or induce superhelical writhe, which in turn can be accommodated through nucleosome remodeling, local alterations in DNA secondary structure, or formation of writhed structures such as plectonemes ^12^.

Although other topological constraints might also be involved, supercoiling-induced plectoneme formation over gene arrays is an intuitive mechanistic explanation for the presence of dinoTADs. An examination of dinoflagellate gene repertoires also corroborates this model, revealing a striking, dinoflagellate-specific expansion of topoisomerase II- and topoisomerase III-like genes (Fig. 1D; Supplementary Fig. 17; Supplementary Table 2), further suggestive of contending with increased levels of writhed forms of helical twist.

Comparison with self-interacting domains in bacteria or *S. pombe* shows much stronger topological insulation for dinoTADs (Supplementary Fig. 14) and 15)). Remarkably, no TAD domains are observed in kinetoplastids, the other lineage with long gene arrays and no transcriptional regulation (Supplementary Fig. 16).

These differences can be rationalized by the unusual dinoflagellate properties. First, neither bacteria nor yeast possess comparably long gene arrays and transcription in those species is highly nonuniform; less transcription-induced torsional stress is therefore expected. Nucleosome loss is the second, and most salient difference. Single mammalian genes as long as dinoTADs are quite common, yet supercoiling is not apparent in mammalian Hi-C maps, nor is it seen in kinetoplastids, which have gene arrays but also have conventional chromatin. We therefore hypothesize that plectonemic structures form due to transcription-induced supercoiling in the nucleosome-depleted genomes of dinoflagellates, while in other eukaryotes, a combination of the wrapping of DNA around nucleosomes, interactions between nucleosomes, and accumulation of DNA twist, prevent their formation (Fig. 2H).

These results generate a number of open questions. How exactly are boundaries between dinoTADs formed mechanistically? Specific boundary elements of markedly different chromatin state could exist; alternatively, these boundaries may self-organize purely through torsion-related mechanisms. The roles that dinoflagellates’ divergent histone genes play is also not clear. Finally, the relationship between Hi-C features and the “toroidal chromonemas” ^13^ observed by electron microscopy remains unknown. Answers to these questions, together with the dissection of specific roles different topoisomerase classes, will help fully elucidate the interplay between packaging proteins, transcription-induced torsional stress, and genome folding in dinoflagellates.

These observations also identify transcription-induced torsional stress as a key direction of future studies in eukaryotes generally. The strength of dinoTADs underlines the potency of this fundamental biological process for generating topological structure. The precise manner by which torsion is accommodated as twist and writhe, as well as its consequences for regulatory protein occupancy, transcriptional activity, and other chromatin processes, such as the behavior of ATP-dependent chromatin remodelers, are exciting questions remaining to be unraveled.

## Author contributions

G.K.M. performed Hi-C experiments. G.K.M and A.E.T. analyzed the data. A.E.T. and T.X. designed and carried out transcription inhibition experiments and cell viability experiments. T.X. carried out *S. minutum* culture and heat stress treatment. W.J.G., A.R.G.. A.K. and J.R.P. supervised the study. G.K.M., A.E.T. and T.X. interpreted data and wrote manuscript with input from all authors.

## Acknowledgments

This work was supported by NIH grants (P50HG007735, RO1 HG008140, U19AI057266 and UM1HG009442 to W.J.G., 1UM1HG009436 to W.J.G. and A.K., 1DP2OD022870-01 and 1U01HG009431 to A.K.), the Rita Allen Foundation (to W.J.G.), the Baxter Foundation Faculty Scholar Grant, and the Human Frontiers Science Program grant RGY006S (to W.J.G). W.J.G is a Chan Zuckerberg Biohub investigator and acknowledges grants 2017-174468 and 2018-182817 from the Chan Zuckerberg Initiative. Fellowship support provided by the Stanford School of Medicine Dean’s Fellowship (G.K.M.), the Siebel Scholars, the Enhancing Diversity in Graduate Education Program and the Weiland Family Fellowship (A.E.T.). This work is also supported by NSF-IOS EDGE Award 1645164 and Carnegie Venture grant 10907 (to T.X. and G.K.M.).

The authors would like to thank Zohar Shipony, Erez Lieberman Aiden, Olga Dudchenko, John R. Pringle, Philip Cleves, and members of the Greenleaf, Kundaje, Pringle and Grossman labs for helpful discussion and suggestions regarding this work.

