## Supplementary Materials for "Transcription-dependent domain-scale 3D genome organization in dinoflagellates"

#### Supplementary Methods

Except where otherwise stated, computational analyses were carried out using custom-written Python scripts.

##### *B. minutum* cell culture

The clonal axenic *Symbiodinium*/*Breviolum minutum* strain SSB01 was used in all experiments. Stock cultures were grown as previously described[14,15] in Daigo's IMK medium for marine microalgae (Wako Pure Chemicals) supplemented with casein hydrolysate (IMK+Cas) at 27 °C at a light intensity of 10 $\mu$mol photons $m^2$ $s^1$ from Philips ALTO II 25-W bulbs on a 12-h-light:12-h-dark cycle. The medium was prepared in artificial seawater (ASW).

##### Transcription inhibition experiments

For $\alpha$-amanitin treatment, *Breviolum minutum* cells at a density of $\sim$1 $\times 10^6$ cells/mL were treated with $\alpha$-amanitin (Sigma-Aldrich, Cat # A2263) at concentrations of 1 $\mu$g/mL ("normal" dose) and 4 $\mu$g/mL ("high") dose.

Samples were harvested at 0, 24, and 48 hours after treatment.

For triptolide treatment, *Breviolum minutum* cells at a density of $\sim$1 $\times 10^6$ cells/mL were treated with triptolide (Sigma-Aldrich, Cat # T3652) at concentrations of 10 $\mu$M ("normal" dose) and 40 $\mu$M ("high") dose.

Samples were harvested fat 0, 8, 24 and 48 hours after treatment.

##### Cell viability measurements

*Photosynthetic activity*

Maximum quantum yields of photosystem II, $Fv/Fm = (Fm - F0)/Fm$ was used to indicate photosynthetic function. S. minutum cultures (approximately $10^6$ cells/mL) were collected and dark adapted for 5 min, and $Fv/Fm$ was determined using a Dual Pam-100 fluorometer (Heinz Walz).

*Colony formation assay*

Fresh SSB01 cells were sampled at 0, 24 and 48 hours after the treatment of transcription inhibitor $\alpha$-amanitin. For each condition, cell suspensions were diluted 1:5 and 1:10 before plating 1 $\mu$L of each dilution on marine broth (BD) agar plates. Plates were incubated at 27 °C at a light intensity of 10 $\mu$mol photons $m^2$ $s^1$. Cell numbers on each plate were counted after three weeks.

##### Hi-C experiments

The in situ Hi-C procedure used to map 3D genomic interactions in *Symbiodinium* was adapted from previous studies[16] as follows:

*Symbiodinium minutum* SSBO1 cells were first crosslinked using 37% formaldehyde (Sigma) at a final concentration of 1% for 15 minutes at room temperature. Formaldehyde was then quenched using 2.5 M Glycine at a final concentration of 0.25 M. Cells were subsequently centrifuged at 2,000 *g* for 5 minutes, washed once in 1$\times$ PBS, and stored at -80 °C.

Cell lysis was initiated by incubation with 250 $\mu$L of cold Hi-C Lysis Buffer (10 mM Tris-HCl pH 8.0, 10 mM NaCl, 0.2% Igepal CA630) on ice for 15 minutes, followed by centrifugation at 2,500 *g* for 5 minutes, a wash with 500 $\mu$L of cold Hi-C Lysis Buffer, and centrifugation at 2,500 *g* for 5 minutes. The pellet was the resuspended in 50 $\mu$L of 0.5% SDS and incubated at 62 °C for 10 minutes. SDS was quenched by adding 145 $\mu$L of $H_2O$ and 25 $\mu$L of 10% Triton X-100 and incubating at 37 °C for 15 minutes.

Restriction digestion was carried out by adding 25 $\mu$L of 10$\times$ NEBuffer 2 and 100 U of the MboI restriction enzyme (NEB, R0147) and incubating for $\geq$2 hours at 37 °C in a Thermomixer at 900 rpm. The reaction was then incubated at 62 °C for 20 minutes in order to inactivate the restriction enzyme.

Fragment ends were filled in by adding 37.5 $\mu$L of 0.4 mM biotin-14-dATP (ThermoFisher Scientific, # 19524-016), 1.5 $\mu$L each of 10 mM dCTP, dGTP and dTTP, and 8 $\mu$L of 5U/$\mu$L DNA Polymerase I Large (Klenow) Fragment (NEB M0210). The reaction was the incubated at 37 °C in a Thermomixer at 900 rpm for 45 minutes.

Fragment end ligation was carried out by adding 663 $\mu$L $H_2O$, 120 $\mu$L 10$\times$ NEB T4 DNA ligase buffer (NEB B0202), 100 $\mu$L of 10% Triton X-100, 12 $\mu$L of 10 mg/mL Bovine Serum Albumin (100$\times$ BSA, NEB), 5 $\mu$L of 400 U/$\mu$L T4 DNA Ligase (NEB M0202), and incubating at room temperature for $\geq$4 hours with rotation.

Nuclei were then pelleted by centrifugation at 3,500 *g* for 5 minutes; the pellet was resuspended in 200 $\mu$L ChIP Elution Buffer (1% SDS, 0.1 M $NaHCO_3$), Proteinase K was added, and incubated at 65 °C overnight to reverse crosslinks.

After addition of 600 $\mu$L 1$\times$TE buffer, DNA was sonicated using a Qsonica S-4000 with a 1/16" tip for 3 minutes, with 10 second pulses at intensity 3.5, and 20 seconds rest between pulses. DNA was then purified using the MinElute PCR Purificaiton Kit (Qiagen #28006), with elution in a total volume of 300 $\mu$L 1$\times$ EB buffer.

For streptavidin pulldown of biotin-labeled DNA, 150

$\mu$L of 10 mg/mL Dynabeads MyOne Streptavidin T1 beads (Life Technologies, 65602) were separated on a magnetic stand, then washed with 400 $\mu$L of 1× TWB (Tween Washing Buffer; 5 mM Tris-HCl pH 7.5; 0.5 mM EDTA; 1 M NaCl; 0.05% Tween 20). The beads were resuspended in 300 $\mu$L of 2× Binding Buffer (10 mM Tris-HCl pH 7.5, 1 mM EDTA; 2 M NaCl), the sonicated DNA was added, and the beads were incubated for ≥15 minutes at room temperature on a rotator. After separation on a magnetic stand, the beads were washed with 600 $\mu$L of 1× TWB, and heated at 55 °C in a Thermomixer with shaking for 2 minutes. After removal of the supernatant on a magnetic stand, the TWB wash and 55 °C incubation were repeated.

Final libraries were prepared on beads using the NEBNext Ultra II DNA Library Prep Kit (NEB, #E7645) as follows. End repair was carried out by resuspending beads in 50 $\mu$L 1× EB buffer, and adding 3 $\mu$L NEB Ultra End Repair Enzyme and 7 $\mu$L NEB Ultra End Repair Enzyme, followed by incubation at 20 °C for 30 minutes and then at 65 °C for 30 minutes.

Adapters were ligated to DNA fragments by adding 30 $\mu$L Blunt Ligation mix, 1 $\mu$L Ligation Enhancer and 2.5 $\mu$L NEB Adapter, incubating at 20 °C for 20 minutes, adding 3 $\mu$L USER enzyme, and incubating at 37 °C for 15 minutes.

Beads were then separated on a magnetic stand, and washed with 600 $\mu$L TWB for 2 minutes at 55 °C, 1000 rpm in a Thermomixer. After separation on a magnetic stand, beads were washed in 100 $\mu$L 0.1 × TE buffer, then resuspended in 16 $\mu$L 0.1 × TE buffer, and heated at 98 °C for 10 minutes.

For PCR, 5 $\mu$L of each of the i5 and i7 NEB Next sequencing adapters were added together with 25 $\mu$L 2× NEB Ultra PCR Mater Mix. PCR was carried out with a 98 °C incubation for 30 seconds and 12 cycles of 98 °C for 10 seconds, 65 °C for 30 seconds, and 72 °C for 1 minute, followed by incubation at 72 °C for 5 minutes.

Beads were separated on a magnetic stand, and the supernatant was cleaned up using 1× AMPure XP beads.

Libraries were sequenced in a paired-end format on a Illumina NextSeq instrument using NextSeq 500/550 high output kits (either 2×75 or 2×36 cycles).

#### Hi-C data processing and assembly scaffolding

As an initial step, Hi-C sequencing reads from all libraries were trimmed of adapter sequences, pooled together, and processed against the previously published *B. minutum* assembly[6] using the Juicer pipeline[17] for analyzing Hi-C datasets (version 1.8.9 of Juicer Tools).

The resulting Hi-C matrices were then used as input to the 3D DNA pipeline[18] for automated scaffolding with the following parameters: `--editor-coarse-resolution 5000 --editor-coarse-region 5000 --polisher-input-size 100000 --polisher-coarse-resolution 1000 --polisher-coarse-region 300000 --splitter-input-size 100000 --splitter-coarse-resolution 5000 --splitter-coarse-region 300000 --sort-output --build-gapped-map -r 10 -i 5000`.

Manual correction of obvious assembly and scaffolding errors was then carried out using Juicebox[17].

After finalizing the scaffolding, Hi-C reads were reprocessed against the new assembly using the Juicer pipeline. This was done individually for each library as well as together for the pooled set of reads.

Data was extracted from the final read matrices using the Juicer suite of tools for Hi-C data analysis.

#### Identification of Hi-C domains

Hi-C matrices were first converted to *cool* format using HiCExplorer[19] "`hicConvertFormat`" with parameters `--inputFormat hic --outputFormat h5` and default resolutions. Subsequent HiCExplorer commands were carried out at 10 kb, 25 kb, and 50 kb resolutions with similar results. Matrices were normalized using "`hicNormalize`" with parameter `--normalize smallest`, and corrected using `''hicCorrectMatrix correct`" with parameters `--correctionMethod KR`. Hi-C domains were computationally identified using the "`hicFindTADs`" from HiCExplorer with parameter `--correctForMultipleTesting fdr`.

#### RNA-seq datasets

Approximately $5 \times 10^7$ cells were collected by centrifugation at 100 *g* for 5 minutes at room temperature. Total RNA was extracted and libraries were constructed for RNA-Seq using the TruSeq RNA Library Prep Kit V2 (Illumina, San Diego, CA, USA) according to the manufacturer protocol. All of the raw sequencing reads are available at Sequence Read Archive (SRA) with accession number SRX7258938.

#### RNA-seq data analysis

RNA-seq reads were aligned against the corresponding assemblies using the STAR aligner[20] (version 2.5.3a) with the following settings: `--limitSjdbInsertNsj 10000000 --outFilterMultimapNmax 50 \verb–`outFilterMismatchNmax— `999 --outFilterMismatchNoverReadLmax 0.04 --alignIntronMin 10 --alignIntronMax 1000000 --alignMatesGapMax 1000000 --alignSJoverhangMin 8 --alignSJDBoverhangMin 1 --sjdbScore 1 --twopassMode Basic --twopass1readsN -1`. As available RNA-seq datasets for *B. minutum* are not strand-specific, the strand orientation of the transcriptome was visualized as follows. Aligned reads were first *de novo* assembled into transcripts and quantified at the transcript level using Stringtie[21] (version 1.3.3.b); the orientation of splice junctions serves as a reliable guide for the directionality of these transcripts. Open reading frames (ORFs) were identified for each transcript, and transcripts with ORFs shorter than 60 amino acids were filtered out of the transcript set. Strand-specific genomic tracks were then generated by assigning to each basepair covered by at least one exon in that set the sum of the TPM (Transcript Per Million transcripts) values of all transcripts it is included in.

##### External Hi-C datasets

Hi-C data for *Trypanosoma brucei* was obtained from GEO accession GSE118764.

Hi-C data for *Schizosaccharomyces pombe* was obtained from GEO accession GSE57316.

Hi-C data for *Caulobacter vibrioides* CB15 was obtained from GEO accession GSE45966.

##### Sequence Analysis

Topoisomerase and other replication-related proteins were identified in annotated MMETSP transcriptome assemblies using HMMER3.0[22] and the Pfam 27.0 protein domain database[23] as previously described[4].

### Supplementary Tables

**Supplementary Table 1:** Summary of Hi-C datasets used in this study

| Hi-C library | Number raw read pairs | Estimated library complexity | Number Hi-C contacts |
|---|---|---|---|
| L142-SSBO1-HIC | 534,609,924 | 920,112,029 | 220,908,462 |
| L533-SSBO1_27C_Hi-C | 556,089,015 | 1,513,268,498 | 151,618,419 |
| L534-SSBO1_34C_Hi-C | 531,461,453 | 2,971,291,849 | 165,231,965 |
| L1240-SSBO1-$\alpha$_amanitin-0h-Hi-C | 111,333,226 | 233,525,989 | 34,384,671 |
| L1241-SSB01-$\alpha$_amanitin-16h-Hi-C-rep1 | 60,696,609 | 317,650,525 | 24,238,281 |
| L1242-SSB01-$\alpha$_amanitin-16h-Hi-C-rep2 | 67,376,168 | 227,736,960 | 25,551,603 |
| L1243-SSB01-$\alpha$_amanitin-24h-Hi-C-rep1 | 81,532,584 | 235,898,386 | 29,748,439 |
| L1244-SSB01-$\alpha$_amanitin-24h-Hi-C-rep2 | 106,381,220 | 110,607,925 | 28,845,306 |
| L1245-SSB01-$\alpha$_amanitin-48h-Hi-C-rep1 | 90,180,763 | 155,046,434 | 27,045,343 |
| L1246-SSB01-$\alpha$_amanitin-48h-Hi-C-rep2 | 78,982,528 | 152,703,652 | 22,153,117 |
| L1247-SSB01-$\alpha$_amanitin_high-48h-Hi-C | 110,015,013 | 157,350,902 | 28,138,017 |
| L1332-SSB01-$\alpha$_amanitin-0h-Hi-C-technical_rep | 117,543,007 | 182,213,300 | 34,089,285 |
| L1333-SSB01-$\alpha$_amanitin-48h-Hi-C-rep1-technical_rep | 117,821,773 | 82,740,021 | 23,654,760 |
| L1334-SSB01-$\alpha$_amanitin_high-48h-Hi-C-technical_rep | 95,662,202 | 164,149,035 | 23,944,231 |
| L1336-SSB01-$\alpha$_amanitin_high-24h-Hi-C-second_time_course | 58,747,402 | 103,174,104 | 15,663,160 |
| L1337-SSB01-$\alpha$_amanitin_high-48h-Hi-C-second_time_course | 83,691,617 | 62,658,394 | 14,523,464 |
| L1344-SSB01-$\alpha$_amanitin/triptolide_0h_NT-Hi-C | 79,383,186 | 208,157,102 | 23,592,335 |
| L1346-SSB01-triptolide_8h_normal_dose-Hi-C | 81,731,190 | 193,514,340 | 22,700,096 |
| L1347-SSB01-triptolide_8h_high_dose-Hi-C | 112,753,865 | 187,235,670 | 28,552,855 |
| L1348-SSB01-Triptolide_24h NT-Hi-C | 52,148,987 | 166,057,825 | 15,674,551 |
| L1349-SSB01-triptolide_24h_normal_dose-Hi-C | 132,715,807 | 206,778,720 | 36,745,591 |
| L1350-SSB01-triptolide_24h_high_dose-Hi-C | 98,429,444 | 265,027,975 | 32,121,298 |
| L1351-SSB01-Triptolide_48h NT-Hi-C | 96,846,551 | 240,797,245 | 28,296,251 |
| L1352-SSB01-triptolide_48h_normal_dose-Hi-C | 85,347,611 | 255,500,603 | 25,051,605 |
| L1353-SSB01-triptolide_48h_high_dose-Hi-C | 99,978,207 | 215,504,692 | 26,572,806 |

**Supplementary Table 2:** Inventory of topoisomerases and some other proteins involved in DNA replication in dinoflagellates and other eukaryotes as annotated by transcriptome assemblies in the MMETSP databses

| clade | species | TOP1 | TOP2 | TOP3 | MCM | PCNA | RPA1 | RPA2 | RPA3 | RFC1 |
|---|---|---|---|---|---|---|---|---|---|---|
| Amoebozoa | *Stereomyxa ramosa* Chinc5 | 1 | 2 | 2 | 6 | 2 | 3 | 0 | 2 | 1 |
| Amoebozoa | *Vexillifera* sp. DIVA3 564 2 | 1 | 2 | 2 | 7 | 1 | 2 | 0 | 0 | 1 |
| Apicomplexa | *Lankesteria abbottii* Grappler Inlet BC | 1 | 1 | 0 | 12 | 5 | 1 | 0 | 0 | 1 |
| Bicosoecid | Bicosoecid sp ms1 | 1 | 0 | 0 | 3 | 1 | 1 | 1 | 1 | 0 |
| Bicosoecid | *Cafeteria roenbergensis* E4 10 | 1 | 0 | 2 | 6 | 1 | 0 | 0 | 1 | 0 |
| Bicosoecid | *Cafeteria* sp. Caron Lab Isolate | 1 | 1 | 4 | 15 | 1 | 1 | 0 | 1 | 1 |
| Bolidophyte | *Bolidomonas pacifica* CCMP 1866 | 2 | 5 | 7 | 8 | 1 | 1 | 0 | 0 | 1 |
| Chlorarachniophyte | *Bigelowiella natans* CCMP1258.1 | 1 | 1 | 9 | 3 | 1 | 4 | 1 | 0 | 0 |
| Chlorarachniophyte | *Bigelowiella natans* CCMP1259 | 1 | 1 | 6 | 7 | 1 | 4 | 1 | 0 | 1 |
| Chlorarachniophyte | *Bigelowiella natans* CCMP 2755 | 0 | 0 | 4 | 5 | 1 | 4 | 1 | 0 | 1 |
| Chlorarachniophyte | *Bigelowiella natans* CCMP623 | 1 | 3 | 7 | 9 | 1 | 2 | 1 | 0 | 1 |
| Chlorarachniophyte | *Chlorarachnion reptans* CCCM449 | 2 | 4 | 8 | 11 | 2 | 3 | 1 | 0 | 1 |
| Chlorarachniophyte | *Lotharella amoebiformis* CCMP2058 | 2 | 6 | 5 | 10 | 1 | 4 | 1 | 0 | 1 |
| Chlorarachniophyte | *Lotharella globosa* CCCM811 | 1 | 2 | 1 | 0 | 1 | 1 | 1 | 1 | 1 |
| Chlorarachniophyte | *Lotharella oceanica* CCMP622 | 1 | 0 | 0 | 1 | 1 | 2 | 1 | 1 | 1 |
| Chlorarachniophyte | *Norrisiella sphaerica* BC52 | 1 | 0 | 3 | 0 | 1 | 2 | 1 | 1 | 0 |
| Chlorarachniophyte | *Partenskyella glossopodia* RCC365 | 1 | 2 | 1 | 7 | 1 | 3 | 1 | 2 | 1 |
| Chlorophyte | *Bathycoccus prasinos* CCMP1898 | 1 | 2 | 3 | 9 | 1 | 2 | 0 | 0 | 0 |
| Chlorophyte | *Bathycoccus prasinos* RCC716 | 1 | 2 | 3 | 7 | 1 | 3 | 0 | 0 | 1 |
| Chlorophyte | *Chlamydomonas* cf sp CCMP681 | 1 | 0 | 0 | 5 | 2 | 1 | 0 | 0 | 1 |
| Chlorophyte | *Crustomastix stigmata* CCMP3273 | 1 | 2 | 4 | 10 | 1 | 1 | 1 | 0 | 1 |
| Chlorophyte | *Cyanoptyche gloeocystis* SAG4.97 | 1 | 0 | 0 | 4 | 1 | 1 | 1 | 0 | 0 |
| Chlorophyte | *Dolichomastix tenuilepis* CCMP3274 | 1 | 1 | 3 | 1 | 2 | 1 | 0 | 1 | 1 |
| Chlorophyte | *Dunaliella tertiolecta* CCMP1320 | 1 | 2 | 3 | 10 | 1 | 2 | 0 | 1 | 1 |
| Chlorophyte | *Mantoniella antarctica* SL 175 | 1 | 8 | 4 | 13 | 1 | 2 | 2 | 1 | 1 |
| Chlorophyte | *Mantoniella* sp CCMP1436 | 1 | 2 | 1 | 2 | 1 | 1 | 1 | 1 | 1 |
| Chlorophyte | *Micromonas* sp CCMP2099 | 1 | 2 | 2 | 9 | 1 | 2 | 0 | 1 | 1 |
| Chlorophyte | *Micromonas* sp NEPCC29 | 1 | 2 | 3 | 7 | 1 | 2 | 0 | 1 | 1 |
| Chlorophyte | *Micromonas* sp RCC472 | 1 | 2 | 2 | 7 | 1 | 2 | 1 | 0 | 1 |
| Chlorophyte | *Nephroselmis pyriformis* CCMP717 | 1 | 4 | 8 | 10 | 1 | 2 | 0 | 1 | 1 |
| Chlorophyte | *Picochlorum oklahomensis* CCMP2329 | 1 | 2 | 2 | 6 | 2 | 2 | 1 | 0 | 1 |
| Chlorophyte | *Picochlorum* sp. RCC944 | 1 | 1 | 2 | 6 | 1 | 2 | 0 | 2 | 1 |
| Chlorophyte | *Picocystis salinarum* CCMP1897 | 1 | 2 | 1 | 8 | 2 | 2 | 1 | 2 | 1 |
| Chlorophyte | *Polytomella parva* SAG 63 3 | 1 | 5 | 3 | 18 | 2 | 3 | 0 | 0 | 1 |
| Chlorophyte | *Prasinoderma coloniale* CCMP1413 | 1 | 2 | 0 | 2 | 1 | 1 | 0 | 0 | 0 |
| Chlorophyte | *Prasinoderma singularis* RCC927 | 1 | 1 | 1 | 7 | 1 | 1 | 0 | 1 | 1 |
| Chlorophyte | *Pterosperma* sp. CCMP1384 | 1 | 0 | 0 | 3 | 1 | 1 | 1 | 1 | 1 |
| Chlorophyte | *Pycnococcus provasolii* RCC2336 | 1 | 1 | 0 | 9 | 1 | 1 | 0 | 0 | 1 |
| Chlorophyte | *Pycnococcus provasolii* RCC931 | 1 | 0 | 0 | 7 | 1 | 1 | 0 | 0 | 1 |
| Chlorophyte | *Pyramimonas parkeae* CCMP726 | 1 | 0 | 4 | 7 | 1 | 2 | 1 | 1 | 1 |
| Chlorophyte | *Stichococcus* sp RCC1054 | 1 | 1 | 1 | 8 | 1 | 1 | 0 | 0 | 1 |
| Chlorophyte | *Tetraselmis chuii* PLY429 | 2 | 0 | 0 | 0 | 0 | 2 | 0 | 1 | 2 |
| Chlorophyte | *Tetraselmis striata* LANL1001 | 1 | 4 | 4 | 11 | 1 | 2 | 0 | 1 | 1 |
| Choanoflagellata | *Acanthoeca* like sp 10tr | 1 | 3 | 4 | 10 | 1 | 1 | 0 | 1 | 1 |
| Chromerida | *Chromera velia* CCMP2878 | 1 | 1 | 3 | 10 | 2 | 2 | 0 | 0 | 1 |
| Chromerida | *Vitrella brassicaformis* CCMP3346 | 1 | 1 | 2 | 9 | 2 | 1 | 0 | 0 | 1 |
| Chrysophyte | *Chromulina nebulosa* UTEXLB2642 | 1 | 1 | 1 | 2 | 1 | 1 | 0 | 0 | 1 |
| Chrysophyte | *Dinobryon* sp UTEXLB2267 | 1 | 3 | 0 | 8 | 1 | 1 | 0 | 0 | 1 |
| Chrysophyte | *Mallomonas* Sp CCMP3275 | 1 | 2 | 1 | 9 | 1 | 1 | 0 | 1 | 1 |
| Chrysophyte | *Ochromonas* sp CCMP1393 | 1 | 2 | 2 | 7 | 1 | 1 | 0 | 0 | 1 |
| Chrysophyte | *Paraphysomonas bandaiensis* Caron Lab Isolate | 1 | 2 | 3 | 9 | 2 | 1 | 1 | 1 | 1 |
| Chrysophyte | *Paraphysomonas imperforata* PA2 | 0 | 1 | 3 | 6 | 1 | 1 | 1 | 1 | 1 |
| Chrysophyte | *Pelagococcus subviridis* CCMP1429 | 1 | 1 | 2 | 11 | 1 | 0 | 0 | 0 | 1 |
| Chrysophyte | *Spumella elongata* CCAP 955 1 | 1 | 1 | 3 | 10 | 4 | 3 | 0 | 1 | 1 |
| Ciliate | *Aristerostoma* sp. ATCC 50986 | 2 | 1 | 1 | 0 | 2 | 1 | 0 | 0 | 2 |
| Ciliate | *Blepharisma japonicum* Stock R1072 | 0 | 0 | 0 | 7 | 4 | 1 | 0 | 0 | 0 |

*Continued on next page*

| clade | species | TOP1 | TOP2 | TOP3 | MCM | PCNA | RPA1 | RPA2 | RPA3 | RFC1 |
|---|---|---|---|---|---|---|---|---|---|---|
| Ciliate | *Climacostomum virens* Stock W 24 | 1 | 2 | 2 | 9 | 3 | 1 | 0 | 0 | 3 |
| Ciliate | *Condylostoma magnum* COL2 | 0 | 0 | 0 | 2 | 0 | 0 | 0 | 0 | 0 |
| Ciliate | *Euplotes focardii* TN1 | 1 | 0 | 0 | 5 | 2 | 1 | 0 | 2 | 0 |
| Ciliate | *Euplotes harpa* FSP1.4 | 2 | 0 | 5 | 3 | 1 | 0 | 0 | 1 | 0 |
| Ciliate | *Fabrea salina* Unknown | 1 | 1 | 3 | 7 | 2 | 3 | 0 | 0 | 2 |
| Ciliate | *Favella taraikaensis* FeNarragansettBay | 0 | 1 | 2 | 7 | 3 | 0 | 0 | 0 | 0 |
| Ciliate | *Litonotus pictus* P1 | 1 | 1 | 2 | 0 | 0 | 0 | 0 | 0 | 0 |
| Ciliate | *Mesodinium pulex* SPMC105 | 2 | 13 | 2 | 16 | 9 | 4 | 0 | 0 | 6 |
| Ciliate | *Myrionecta rubra* CCMP2563 | 0 | 1 | 4 | 11 | 1 | 1 | 0 | 1 | 0 |
| Ciliate | *Platyophrya macrostoma* WH | 4 | 4 | 4 | 23 | 4 | 6 | 0 | 0 | 3 |
| Ciliate | *Protocruzia adherens* Boccale | 3 | 1 | 0 | 9 | 3 | 3 | 0 | 0 | 1 |
| Ciliate | *Pseudokeronopsis* sp. OXSARD2 | 1 | 1 | 1 | 6 | 1 | 0 | 0 | 1 | 1 |
| Ciliate | *Strombidinopsis acuminatum* SPMC142 | 2 | 6 | 0 | 32 | 10 | 5 | 0 | 0 | 0 |
| Ciliate | *Strombidinopsis* sp. SopsisLIS2011 | 1 | 0 | 0 | 8 | 3 | 2 | 0 | 0 | 0 |
| Ciliate | *Strombidium* inclinatum S3 | 1 | 1 | 2 | 8 | 1 | 1 | 0 | 0 | 1 |
| Ciliate | *Strombidium rassoulzadegani* ras09 | 1 | 0 | 1 | 6 | 1 | 1 | 0 | 1 | 0 |
| Ciliate | *Tiarina fusus* LIS | 1 | 7 | 3 | 16 | 3 | 4 | 2 | 1 | 1 |
| Cryptophyte | *Chroomonas mesostigmatica* cf CCMP1168 | 1 | 5 | 4 | 8 | 1 | 2 | 2 | 0 | 1 |
| Cryptophyte | *Cryptomonas curvata* CCAP979 52 | 2 | 0 | 2 | 0 | 1 | 1 | 0 | 1 | 0 |
| Cryptophyte | *Cryptomonas paramecium* CCAP977 2a | 3 | 2 | 2 | 5 | 1 | 1 | 0 | 0 | 1 |
| Cryptophyte | *Geminigera cryophila* CCMP2564 | 2 | 1 | 5 | 11 | 1 | 2 | 0 | 1 | 2 |
| Cryptophyte | *Geminigera* sp. Caron Lab Isolate | 1 | 3 | 5 | 18 | 1 | 5 | 0 | 1 | 1 |
| Cryptophyte | *Goniomonas pacifica* CCMP1869 | 8 | 4 | 4 | 12 | 1 | 5 | 1 | 3 | 7 |
| Cryptophyte | *Guillardia theta* CCMP 2712 | 1 | 0 | 2 | 3 | 1 | 1 | 0 | 1 | 0 |
| Cryptophyte | *Hemiselmis andersenii* CCMP644 | 1 | 2 | 5 | 12 | 1 | 2 | 0 | 1 | 1 |
| Cryptophyte | *Hemiselmis rufescens* PCC563 | 1 | 0 | 3 | 7 | 1 | 1 | 1 | 1 | 1 |
| Cryptophyte | *Hemiselmis tepida* CCMP443 | 3 | 2 | 0 | 3 | 1 | 1 | 1 | 1 | 1 |
| Cryptophyte | *Hemiselmis viresens* PCC157 | 1 | 0 | 0 | 7 | 1 | 1 | 0 | 1 | 0 |
| Cryptophyte | *Palpitomonas bilix* NIES 2562 | 0 | 1 | 2 | 13 | 4 | 3 | 0 | 1 | 3 |
| Cryptophyte | *Proteomonas sulcata* CCMP704 | 0 | 1 | 0 | 3 | 1 | 1 | 0 | 0 | 1 |
| Cryptophyte | *Rhodomonas lens* RHODO | 2 | 3 | 2 | 2 | 2 | 2 | 0 | 1 | 0 |
| Cryptophyte | *Rhodomonas* sp. CCMP768 | 1 | 0 | 1 | 0 | 1 | 1 | 0 | 0 | 0 |
| Diatome | *Amphiprora* sp. | 1 | 4 | 3 | 9 | 1 | 1 | 0 | 0 | 1 |
| Diatome | *Amphora coffeaeformis* CCMP127 | 1 | 1 | 0 | 4 | 1 | 1 | 0 | 0 | 0 |
| Diatome | *Asterionellopsis glacialis* CCMP134 | 1 | 7 | 1 | 10 | 1 | 1 | 0 | 0 | 1 |
| Diatome | *Astrosyne radiata* 13vi08 1A | 1 | 8 | 3 | 6 | 3 | 2 | 0 | 0 | 1 |
| Diatome | *Attheya septentrionalis* CCMP2084 | 1 | 2 | 0 | 9 | 1 | 1 | 0 | 0 | 1 |
| Diatome | *Aulacoseira subarctica* CCAP 1002 5 | 1 | 2 | 3 | 8 | 2 | 1 | 0 | 0 | 1 |
| Diatome | *Chaetoceros affinis* CCMP159 | 1 | 3 | 1 | 8 | 1 | 1 | 0 | 0 | 1 |
| Diatome | *Chaetoceros curvisetus* | 1 | 4 | 4 | 6 | 1 | 3 | 0 | 0 | 1 |
| Diatome | *Chaetoceros debilis* MM31A_1 | 1 | 3 | 1 | 12 | 1 | 1 | 0 | 0 | 1 |
| Diatome | *Chaetoceros neogracile* CCMP1317 | 1 | 9 | 3 | 10 | 1 | 1 | 0 | 1 | 1 |
| Diatome | *Coscinodiscus wailesii* CCMP2513 | 1 | 3 | 6 | 10 | 1 | 1 | 0 | 1 | 1 |
| Diatome | *Craspedostauros australis* CCMP3328 | 1 | 0 | 0 | 4 | 0 | 1 | 0 | 0 | 0 |
| Diatome | *Cyclophora tenuis* ECT3854 | 1 | 1 | 0 | 3 | 1 | 1 | 0 | 0 | 0 |
| Diatome | *Cyclotella meneghiniana* CCMP 338 | 1 | 4 | 3 | 8 | 1 | 1 | 0 | 0 | 1 |
| Diatome | *Cylindrotheca closterium* KMMCC:B 181 | 3 | 7 | 3 | 14 | 1 | 2 | 0 | 0 | 1 |
| Diatome | *Dactyliosolen fragilissimus* Unknown | 1 | 3 | 3 | 8 | 1 | 1 | 0 | 1 | 1 |
| Diatome | *Ditylum brightwellii* GSO103 | 1 | 4 | 3 | 11 | 1 | 1 | 0 | 1 | 1 |
| Diatome | *Ditylum brightwellii* GSO104 | 1 | 4 | 5 | 10 | 1 | 1 | 0 | 1 | 1 |
| Diatome | *Ditylum brightwellii* GSO105 | 1 | 2 | 3 | 11 | 2 | 1 | 0 | 1 | 1 |
| Diatome | *Entomoneis* sp. CCMP2396 | 0 | 1 | 0 | 0 | 1 | 1 | 0 | 0 | 0 |
| Diatome | *Eucampia antarctica* CCMP1452 | 1 | 3 | 0 | 5 | 1 | 1 | 1 | 1 | 1 |
| Diatome | *Extubocellulus spinifer* CCMP396 | 1 | 4 | 10 | 13 | 2 | 5 | 3 | 1 | 2 |
| Diatome | *Fragilariopsis kerguelensis* L2_C3 | 1 | 3 | 3 | 11 | 1 | 1 | 2 | 0 | 1 |
| Diatome | *Fragilariopsis kerguelensis* L26_C5 | 1 | 3 | 5 | 22 | 1 | 1 | 3 | 0 | 1 |
| Diatome | *Grammatophora oceanica* CCMP 410 | 1 | 1 | 3 | 5 | 1 | 1 | 0 | 0 | 1 |
| Diatome | *Helicotheca tamensis* CCMP826 | 0 | 1 | 0 | 1 | 1 | 1 | 0 | 1 | 0 |
| Diatome | *Leptocylindrus danicus* var. apora B651 | 3 | 5 | 3 | 0 | 3 | 2 | 0 | 1 | 1 |

*Continued on next page*

| clade | species | TOP1 | TOP2 | TOP3 | MCM | PCNA | RPA1 | RPA2 | RPA3 | RFC1 |
|---|---|---|---|---|---|---|---|---|---|---|
| Diatome | *Leptocylindrus danicus* var. danicus B650 | 3 | 11 | 3 | 19 | 1 | 1 | 0 | 1 | 2 |
| Diatome | *Licmophora paradoxa* CCMP2313 | 1 | 1 | 3 | 7 | 1 | 2 | 0 | 0 | 1 |
| Diatome | *Minutocellus polymorphus* CCMP3303 | 0 | 0 | 0 | 3 | 1 | 1 | 1 | 1 | 0 |
| Diatome | *Minutocellus polymorphus* NH13 | 2 | 8 | 7 | 21 | 1 | 0 | 1 | 0 | 3 |
| Diatome | *Minutocellus polymorphus* RCC2270 | 1 | 2 | 1 | 7 | 1 | 1 | 1 | 1 | 1 |
| Diatome | *Nitzschia punctata* CCMP561 | 1 | 2 | 2 | 9 | 1 | 1 | 1 | 1 | 1 |
| Diatome | *Odontella aurita* isolate 1302 5 | 1 | 3 | 7 | 11 | 2 | 2 | 1 | 1 | 1 |
| Diatome | *Odontella sinensis* Grunow 1884 | 1 | 3 | 0 | 2 | 1 | 1 | 1 | 1 | 1 |
| Diatome | *Proboscia alata* PI_D3 | 1 | 7 | 2 | 21 | 1 | 1 | 2 | 0 | 1 |
| Diatome | *Pseudo-nitzschia australis* 10249_10_AB | 1 | 3 | 4 | 8 | 1 | 1 | 1 | 0 | 1 |
| Diatome | *Pseudo-nitzschia fradulenta* WWA7 | 2 | 11 | 6 | 24 | 4 | 5 | 0 | 0 | 3 |
| Diatome | *Rhizosolenia setigera* CCMP 1694 | 1 | 7 | 4 | 18 | 1 | 2 | 0 | 0 | 2 |
| Diatome | *Skeletonema dohrnii* SkelB | 1 | 2 | 0 | 14 | 1 | 1 | 2 | 1 | 1 |
| Diatome | *Skeletonema marinoi* SkelA | 1 | 1 | 2 | 7 | 1 | 1 | 2 | 0 | 1 |
| Diatome | *Skeletonema menzelii* CCMP793 | 1 | 4 | 4 | 8 | 1 | 1 | 2 | 0 | 1 |
| Diatome | *Stauroneis constricta* CCMP1120 | 1 | 0 | 1 | 1 | 1 | 1 | 1 | 0 | 0 |
| Diatome | *Staurosira complex* sp. CCMP2646 | 1 | 3 | 4 | 8 | 1 | 1 | 0 | 1 | 1 |
| Diatome | *Stephanopyxis turris* CCMP 815 | 2 | 0 | 1 | 7 | 3 | 2 | 0 | 1 | 1 |
| Diatome | *Striatella unipunctata* CCMP2910 | 4 | 2 | 1 | 6 | 3 | 0 | 1 | 0 | 2 |
| Diatome | *Synedropsis recta* cf CCMP1620 | 1 | 2 | 0 | 1 | 1 | 1 | 1 | 1 | 0 |
| Diatome | *Thalassionema frauenfeldii* CCMP 1798 | 1 | 5 | 7 | 15 | 1 | 3 | 1 | 1 | 2 |
| Diatome | *Thalassionema nitzschioides* L26_B | 1 | 3 | 4 | 8 | 1 | 1 | 1 | 1 | 1 |
| Diatome | *Thalassiosira antarctica* CCMP982 | 1 | 4 | 2 | 12 | 1 | 1 | 3 | 1 | 1 |
| Diatome | *Thalassiosira gravida* GMp14c1 | 1 | 1 | 3 | 13 | 1 | 1 | 2 | 1 | 1 |
| Diatome | *Thalassiosira miniscula* CCMP1093 | 1 | 13 | 6 | 10 | 1 | 1 | 2 | 1 | 1 |
| Diatome | *Thalassiosira oceanica* CCMP1005 | 1 | 10 | 1 | 10 | 1 | 1 | 0 | 0 | 1 |
| Diatome | *Thalassiosira rotula* CCMP3096 | 1 | 5 | 3 | 11 | 1 | 1 | 2 | 1 | 1 |
| Diatome | *Thalassiosira rotula* GSO102 | 1 | 3 | 2 | 11 | 1 | 1 | 1 | 1 | 1 |
| Diatome | *Thalassiosira weissflogii* CCMP1010 | 1 | 4 | 1 | 9 | 1 | 0 | 1 | 0 | 1 |
| Diatome | *Thalassiosira weissflogii* CCMP1336 | 1 | 4 | 1 | 8 | 1 | 0 | 1 | 0 | 1 |
| Diatome | *Thalassiothrix antarctica* L6_D1 | 1 | 2 | 4 | 6 | 1 | 1 | 0 | 1 | 1 |
| Diatome | *Triceratium dubium* CCMP147 | 0 | 1 | 1 | 1 | 1 | 0 | 1 | 1 | 0 |
| Dinoflagellata | *Alexandrium temarense* CCMP1771 | 3 | 18 | 12 | 45 | 18 | 10 | 3 | 4 | 2 |
| Dinoflagellata | *Amphidinium carterae* CCMP1314 | 2 | 5 | 5 | 8 | 2 | 4 | 0 | 0 | 3 |
| Dinoflagellata | *Azadinium spinosum* 3D9 | 1 | 12 | 13 | 35 | 11 | 6 | 0 | 0 | 3 |
| Dinoflagellata | *Brandtodinium nutriculum* RCC3387 | 1 | 13 | 9 | 30 | 21 | 4 | 0 | 0 | 3 |
| Dinoflagellata | *Ceratium fusus* PA161109 | 1 | 15 | 10 | 18 | 12 | 9 | 1 | 1 | 3 |
| Dinoflagellata | *Crypthecodinium cohnii* Seligo | 1 | 6 | 5 | 15 | 2 | 4 | 0 | 0 | 3 |
| Dinoflagellata | *Dinophysis acuminata* DAEP01 | 4 | 15 | 9 | 29 | 13 | 8 | 0 | 0 | 2 |
| Dinoflagellata | *Durinskia baltica* CSIRO_CS 38 | 2 | 12 | 9 | 18 | 9 | 8 | 0 | 0 | 4 |
| Dinoflagellata | *Gambierdiscus australes* CAWD 149 | 1 | 5 | 0 | 9 | 14 | 6 | 0 | 0 | 2 |
| Dinoflagellata | *Glenodinium foliaceum* CCAP1116_3 | 2 | 9 | 3 | 23 | 7 | 6 | 0 | 1 | 4 |
| Dinoflagellata | *Gonyaulax spinifera* CCMP409 | 1 | 2 | 0 | 10 | 10 | 8 | 1 | 1 | 1 |
| Dinoflagellata | *Heterocapsa rotundata* SCCAP K 0483 | 2 | 19 | 4 | 12 | 6 | 4 | 0 | 0 | 6 |
| Dinoflagellata | *Heterocapsa triquestra* CCMP 448 | 1 | 8 | 5 | 13 | 5 | 4 | 0 | 0 | 3 |
| Dinoflagellata | *Karenia brevis* CCMP2229 | 1 | 14 | 8 | 10 | 8 | 7 | 0 | 1 | 4 |
| Dinoflagellata | *Karenia brevis* SP1 | 1 | 14 | 13 | 16 | 6 | 8 | 0 | 1 | 4 |
| Dinoflagellata | *Karenia brevis* SP3 | 1 | 12 | 9 | 13 | 8 | 10 | 0 | 1 | 4 |
| Dinoflagellata | *Karenia brevis* Wilson | 1 | 14 | 7 | 14 | 9 | 8 | 0 | 2 | 5 |
| Dinoflagellata | *Karlodinium micrum* CCMP2283 | 2 | 9 | 7 | 46 | 13 | 31 | 2 | 0 | 5 |
| Dinoflagellata | *Kryptoperidinium foliaceum* CCMP1326 | 4 | 14 | 11 | 64 | 16 | 10 | 1 | 0 | 7 |
| Dinoflagellata | *Lingulodinium polyedra* CCMP1738 | 1 | 17 | 8 | 19 | 11 | 11 | 1 | 0 | 3 |
| Dinoflagellata | *Noctiluca scintillans* Unknown | 1 | 7 | 3 | 9 | 1 | 6 | 0 | 1 | 1 |
| Dinoflagellata | *Oxyrrhis marina* | 1 | 2 | 5 | 9 | 7 | 3 | 0 | 1 | 2 |
| Dinoflagellata | *Oxyrrhis marina* CCMP1795 | 0 | 0 | 0 | 0 | 3 | 0 | 0 | 0 | 0 |
| Dinoflagellata | *Oxyrrhis marina* LB1974 | 1 | 2 | 4 | 10 | 4 | 2 | 0 | 0 | 2 |
| Dinoflagellata | *Pelagodinium beii* RCC1491 | 1 | 8 | 2 | 12 | 11 | 4 | 0 | 0 | 4 |
| Dinoflagellata | *Peridinium aciculiferum* PAER_2 | 1 | 7 | 5 | 11 | 6 | 5 | 0 | 0 | 3 |
| Dinoflagellata | *Polarella glacialis* CCMP 1383 | 1 | 28 | 5 | 23 | 5 | 5 | 0 | 0 | 8 |

*Continued on next page*

Supplementary Table 2 – *Continued from previous page*

| clade | species | TOP1 | TOP2 | TOP3 | MCM | PCNA | RPA1 | RPA2 | RPA3 | RFC1 |
|---|---|---|---|---|---|---|---|---|---|---|
| Dinoflagellata | *Prorocentrum minimum* CCMP1329 | 1 | 15 | 6 | 29 | 13 | 6 | 0 | 0 | 3 |
| Dinoflagellata | *Prorocentrum minimum* CCMP2233 | 1 | 14 | 4 | 29 | 12 | 5 | 0 | 0 | 3 |
| Dinoflagellata | *Protoceratium reticulatum* CCCM 535 CCMP 1889 | 2 | 20 | 9 | 18 | 11 | 10 | 0 | 0 | 2 |
| Dinoflagellata | *Pyrodinium bahamense* pbaha01 | 1 | 21 | 8 | 29 | 19 | 11 | 0 | 0 | 3 |
| Dinoflagellata | *Scrippsiella hangoei* like SHHI_4 | 1 | 8 | 6 | 22 | 6 | 16 | 0 | 2 | 2 |
| Dinoflagellata | *Scrippsiella hangoei* SHTV5 | 1 | 8 | 11 | 14 | 3 | 5 | 0 | 0 | 2 |
| Dinoflagellata | *Scrippsiella trochoidea* CCMP3099 | 1 | 27 | 10 | 38 | 12 | 8 | 1 | 1 | 3 |
| Dinoflagellata | *Symbiodinium kawagutii* CCMP2468 | 0 | 0 | 0 | 0 | 2 | 0 | 0 | 0 | 0 |
| Dinoflagellata | *Symbiodinium* sp. C1 | 1 | 9 | 4 | 9 | 6 | 4 | 0 | 0 | 3 |
| Dinoflagellata | *Symbiodinium* sp. C15 | 1 | 7 | 2 | 12 | 3 | 4 | 1 | 0 | 3 |
| Dinoflagellata | *Symbiodinium* sp. CCMP2430 | 1 | 7 | 2 | 10 | 7 | 4 | 0 | 0 | 3 |
| Dinoflagellata | *Symbiodinium* sp. Mp | 1 | 7 | 3 | 13 | 6 | 3 | 0 | 0 | 3 |
| Dinoflagellata | *Togula jolla* CCCM 725 | 1 | 17 | 3 | 21 | 3 | 6 | 0 | 0 | 4 |
| Discosea | *Mayorella* sp. BSH 02190019 | 1 | 3 | 2 | 5 | 1 | 1 | 0 | 1 | 1 |
| Discosea | *Neoparamoeba aestuarina* SoJaBio B1 5 56 2 | 3 | 3 | 3 | 12 | 3 | 3 | 0 | 1 | 1 |
| Discosea | *Paramoeba atlantica* 621 1 CCAP 1560 9 | 1 | 3 | 2 | 8 | 3 | 2 | 0 | 1 | 1 |
| Discosea | *Pessonella* sp. PRA 29 | 1 | 1 | 3 | 0 | 1 | 5 | 0 | 2 | 3 |
| Discosea | *Stygamoeba regulata* BSH 02190019 | 3 | 8 | 2 | 7 | 2 | 4 | 0 | 0 | 2 |
| Discosea | *Trichosphaerium* sp. Am I 7 wt | 2 | 0 | 0 | 1 | 2 | 3 | 0 | 0 | 2 |
| Eugelnophyta | *Eutreptiella gymnastica* like CCMP1594 | 1 | 1 | 1 | 5 | 2 | 1 | 0 | 1 | 1 |
| Foraminifera | *Ammonia* sp. Unknown | 1 | 1 | 3 | 9 | 5 | 2 | 0 | 1 | 1 |
| Foraminifera | *Elphidium margaritaceum* Unknown | 1 | 1 | 2 | 8 | 3 | 1 | 1 | 0 | 1 |
| Foraminifera | *Rosalina* sp. Unknown | 1 | 0 | 0 | 9 | 5 | 0 | 2 | 0 | 1 |
| Foraminifera | *Sorites* sp. Unknown | 3 | 3 | 0 | 27 | 12 | 3 | 0 | 0 | 2 |
| Fungi | *Debaryomyces hansenii* J26 | 1 | 0 | 0 | 4 | 0 | 0 | 0 | 0 | 1 |
| Glaucophyte | *Gloeochaete witrockiana* SAG46_84 | 2 | 2 | 3 | 9 | 2 | 2 | 1 | 1 | 1 |
| Haptophyte | *Calcidiscus leptoporus* RCC1130 | 1 | 3 | 0 | 7 | 1 | 1 | 1 | 0 | 1 |
| Haptophyte | *Chrysochromulina brevifilum* UTEX LB 985 | 1 | 2 | 1 | 4 | 1 | 3 | 0 | 1 | 0 |
| Haptophyte | *Chrysochromulina ericina* CCMP281 | 2 | 1 | 0 | 10 | 1 | 3 | 1 | 1 | 2 |
| Haptophyte | *Chrysochromulina polylepis* CCMP1757 | 1 | 3 | 5 | 9 | 1 | 2 | 1 | 1 | 1 |
| Haptophyte | *Chrysoculter rhomboideus* RCC1486 | 1 | 0 | 0 | 9 | 1 | 0 | 0 | 1 | 0 |
| Haptophyte | *Coccolithus pelagicus* ssp *braarudi* PLY182g | 1 | 3 | 0 | 7 | 1 | 2 | 1 | 1 | 0 |
| Haptophyte | *Emiliania huxleyi* 374 | 1 | 2 | 1 | 9 | 1 | 1 | 0 | 0 | 0 |
| Haptophyte | *Emiliania huxleyi* 379 | 1 | 1 | 1 | 0 | 0 | 2 | 0 | 0 | 0 |
| Haptophyte | *Emiliania huxleyi* CCMP370 | 1 | 3 | 5 | 9 | 0 | 2 | 1 | 1 | 1 |
| Haptophyte | *Emiliania huxleyi* PLYM219 | 1 | 3 | 4 | 10 | 0 | 2 | 1 | 1 | 1 |
| Haptophyte | *Exanthemachrysis gayraliae* RCC1523 | 1 | 2 | 0 | 1 | 1 | 1 | 0 | 1 | 1 |
| Haptophyte | *Gephyrocapsa oceanica* RCC1303 | 1 | 3 | 5 | 11 | 1 | 1 | 0 | 0 | 1 |
| Haptophyte | *Imantonia* sp. RCC918 | 3 | 1 | 1 | 4 | 2 | 1 | 1 | 1 | 0 |
| Haptophyte | *Isochrysis galbana* CCMP1323 | 2 | 5 | 6 | 13 | 2 | 3 | 1 | 0 | 2 |
| Haptophyte | *Isochrysis* sp. CCMP1244 | 1 | 2 | 5 | 11 | 1 | 1 | 0 | 1 | 1 |
| Haptophyte | *Isochrysis* sp. CCMP1324 | 1 | 2 | 0 | 12 | 1 | 2 | 1 | 0 | 1 |
| Haptophyte | *Pavlova* sp. CCMP459 | 1 | 2 | 1 | 6 | 2 | 1 | 2 | 1 | 1 |
| Haptophyte | *Phaeocystis antarctica* Caron Lab Isolate | 3 | 7 | 2 | 12 | 1 | 3 | 2 | 0 | 2 |
| Haptophyte | *Phaeocystis* sp. CCMP2710 | 1 | 0 | 1 | 2 | 1 | 1 | 1 | 1 | 1 |
| Haptophyte | *Pleurochrysis carterae* CCMP645 | 3 | 2 | 1 | 7 | 1 | 2 | 1 | 1 | 1 |
| Haptophyte | *Prymnesium parvum* Texoma1 | 1 | 6 | 4 | 1 | 1 | 2 | 1 | 1 | 1 |
| Haptophyte | *Scyphosphaera apsteinii* RCC1455 | 1 | 3 | 1 | 7 | 1 | 2 | 1 | 0 | 1 |
| Heterolobosea | *Percolomonas cosmopolitus* AE 1 ATCC 50343 | 1 | 4 | 2 | 9 | 2 | 2 | 0 | 0 | 1 |
| Heterolobosea | *Percolomonas cosmopolitus* WS | 1 | 3 | 1 | 12 | 1 | 2 | 0 | 0 | 3 |
| Khakista | *Corethron pennatum* L29A3 | 2 | 5 | 5 | 16 | 1 | 1 | 1 | 0 | 1 |
| Khakista | *Detonula confervacea* CCMP 353 | 1 | 3 | 2 | 9 | 1 | 1 | 2 | 1 | 1 |
| Kinetoplastida | *Neobodo designis* CCAP 1951 1 | 1 | 1 | 4 | 8 | 1 | 1 | 0 | 0 | 1 |
| Labyrinthulida | *Aplanochytrium* sp. PBS07 | 1 | 2 | 1 | 3 | 1 | 1 | 1 | 2 | 1 |
| Labyrinthulida | *Aplanochytrium stocchinoi* GSBS06 | 1 | 2 | 0 | 7 | 1 | 1 | 1 | 1 | 1 |
| Pelagophyte | *Aureococcus anophagefferens* CCMP1850 | 6 | 2 | 3 | 45 | 1 | 2 | 0 | 0 | 1 |
| Pelagophyte | *Aureoumbra lagunensis* CCMP1510 | 1 | 2 | 2 | 9 | 1 | 2 | 1 | 0 | 1 |
| Pelagophyte | *Chrysocystis fragilis* CCMP3189 | 2 | 0 | 2 | 6 | 1 | 1 | 1 | 0 | 1 |

*Continued on next page*

Supplementary Table 2 – *Continued from previous page*

| clade | species | TOP1 | TOP2 | TOP3 | MCM | PCNA | RPA1 | RPA2 | RPA3 | RFC1 |
|---|---|---|---|---|---|---|---|---|---|---|
| Pelagophyte | *Chrysoreinhardia* sp. CCMP2950 | 1 | 2 | 1 | 5 | 0 | 1 | 0 | 0 | 1 |
| Pelagophyte | *Chrysoreinhardia* sp. CCMP3193 | 1 | 3 | 2 | 10 | 1 | 2 | 1 | 0 | 1 |
| Pelagophyte | *Pelagomonas calceolata* CCMP1756 | 1 | 2 | 1 | 9 | 1 | 1 | 2 | 0 | 1 |
| Pelagophyte | *Sarcinochrysis* sp. CCMP770 | 0 | 0 | 0 | 2 | 1 | 1 | 1 | 1 | 0 |
| Perkinsid | *Perkinsus chesapeaki* ATCC_PRA_65 | 2 | 0 | 0 | 0 | 0 | 0 | 0 | 0 | 0 |
| Perkinsid | *Perkinsus marinus* ATCC50439 | 1 | 0 | 0 | 1 | 2 | 0 | 0 | 0 | 0 |
| Pinguiophyte | *Phaeomonas parva* CCMP2877 | 1 | 3 | 1 | 5 | 3 | 1 | 0 | 1 | 0 |
| Pinguiophyte | *Pinguiococcus pyrenoidosus* CCMP2078 | 1 | 2 | 3 | 0 | 1 | 1 | 0 | 1 | 0 |
| Raphidophyte | *Chattonella subsalsa* CCMP2191 | 1 | 3 | 0 | 5 | 1 | 1 | 1 | 0 | 1 |
| Raphidophyte | *Fibrocapsa japonica* CCMP1661 | 0 | 1 | 1 | 5 | 1 | 1 | 0 | 1 | 0 |
| Raphidophyte | *Heterosigma akashiwo* CCMP2393 | 1 | 4 | 2 | 11 | 2 | 1 | 0 | 1 | 1 |
| Raphidophyte | *Heterosigma akashiwo* CCMP3107 | 1 | 7 | 2 | 0 | 1 | 1 | 0 | 0 | 0 |
| Raphidophyte | *Heterosigma akashiwo* CCMP452 | 0 | 1 | 0 | 4 | 1 | 1 | 0 | 0 | 0 |
| Raphidophyte | *Heterosigma akashiwo* NB | 1 | 6 | 1 | 8 | 1 | 1 | 0 | 1 | 1 |
| Rhodophyte | *Compsopogon coeruleus* SAG 36.94 | 1 | 3 | 2 | 11 | 1 | 1 | 0 | 0 | 1 |
| Rhodophyte | *Erythrolobus australicus* CCMP3124 | 1 | 2 | 3 | 0 | 1 | 1 | 0 | 1 | 1 |
| Rhodophyte | *Erythrolobus madagascarensis* CCMP3276 | 1 | 1 | 1 | 3 | 1 | 2 | 0 | 1 | 0 |
| Rhodophyte | *Madagascaria erythrocladiodes* CCMP3234 | 3 | 4 | 5 | 12 | 1 | 2 | 0 | 1 | 2 |
| Rhodophyte | *Porphyridium aerugineum* SAG 1380 2 | 2 | 1 | 2 | 5 | 1 | 2 | 1 | 0 | 1 |
| Rhodophyte | *Rhodella maculata* CCMP736 | 1 | 3 | 3 | 12 | 1 | 1 | 0 | 0 | 1 |
| Rhodophyte | *Rhodosorus marinus* CCMP 769 | 1 | 8 | 6 | 17 | 0 | 3 | 0 | 0 | 2 |
| Rhodophyte | *Timspurckia oligopyrenoides* CCMP3278 | 1 | 2 | 4 | 6 | 1 | 2 | 1 | 1 | 1 |
| Silicoflagellates | *Dictyocha speculum* CCMP1381 | 1 | 4 | 2 | 9 | 1 | 2 | 1 | 1 | 1 |
| Silicoflagellates | *Pseudopedinella elastica* CCMP716 | 1 | 5 | 6 | 9 | 1 | 1 | 1 | 1 | 1 |
| Silicoflagellates | *Pteridomonas danica* PT | 1 | 1 | 1 | 2 | 1 | 1 | 1 | 1 | 0 |
| Silicoflagellates | *Rhizochromulina marina* cf CCMP1243 | 1 | 5 | 2 | 8 | 2 | 2 | 1 | 1 | 1 |
| Synchromophyceae | *Synchroma pusillum* CCMP3072 | 1 | 0 | 1 | 3 | 3 | 1 | 0 | 1 | 1 |
| Syndinian | *Amoebophrya* sp. Ameob2 | 2 | 8 | 1 | 13 | 0 | 1 | 0 | 0 | 0 |
| Thraustochytrid | *Aurantiochytrium limacinum* ATCCMYA1381 | 1 | 3 | 2 | 9 | 1 | 1 | 0 | 1 | 1 |
| Thraustochytrid | *Schizochytrium aggregatum* ATCC28209 | 1 | 1 | 1 | 4 | 1 | 1 | 0 | 0 | 1 |
| Thraustochytrid | *Thraustochytrium* sp. LLF1b | 1 | 2 | 1 | 9 | 1 | 1 | 0 | 1 | 1 |
| Tubulinid | *Filamoeba nolandi* NC AS 23 1 | 2 | 4 | 1 | 13 | 0 | 3 | 1 | 0 | 1 |
| Tubulinid | *Sexangularia* sp. ATCC50979 | 0 | 6 | 7 | 14 | 2 | 2 | 1 | 0 | 3 |
| Vanellinid | *Vannella robusta* DIVA3 518 3 11 1 6 | 1 | 2 | 3 | 6 | 1 | 2 | 1 | 1 | 1 |
| Vanellinid | *Vannella* sp. DIVA3 517 6 12 | 6 | 6 | 9 | 13 | 1 | 1 | 0 | 1 | 1 |
| Xantophyte | *Vaucheria litorea* CCMP2940 | 1 | 2 | 0 | 6 | 1 | 1 | 0 | 1 | 1 |

#### Supplementary Figures

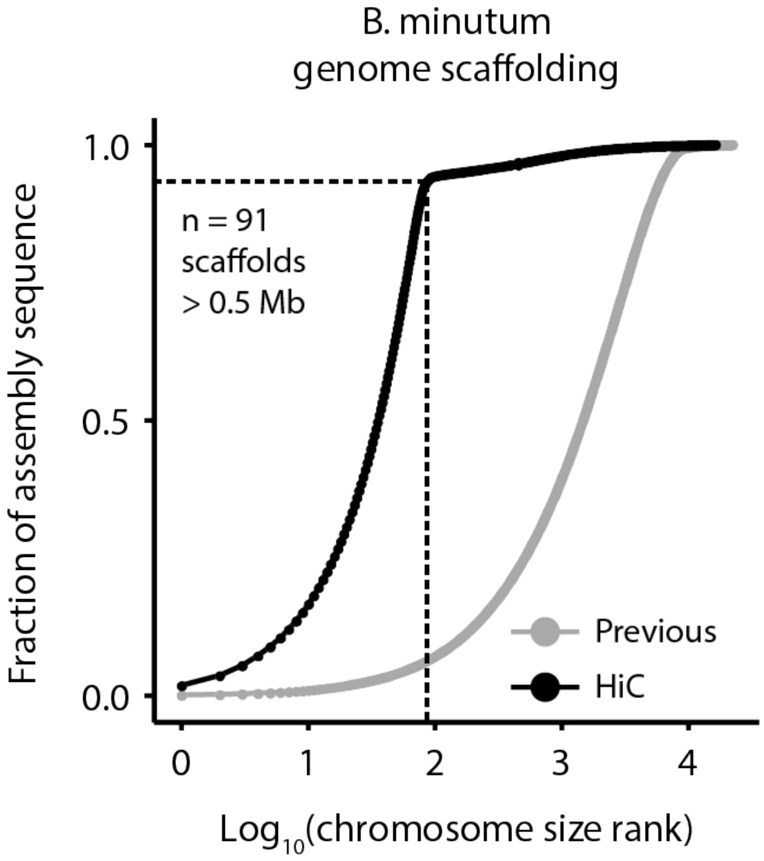


**Supplementary Figure 1: Cumulative distribution of scaffolds and pseudochromosome sizes before and after Hi-C scaffolding of the draft *Breviolum/Symbiodinium minutum* assembly[6].** 3D DNA[18] scaffolding of the assembly results in 91 major pseudochromosomes ≥500kb encompassing ∼94% of the assembled sequence.

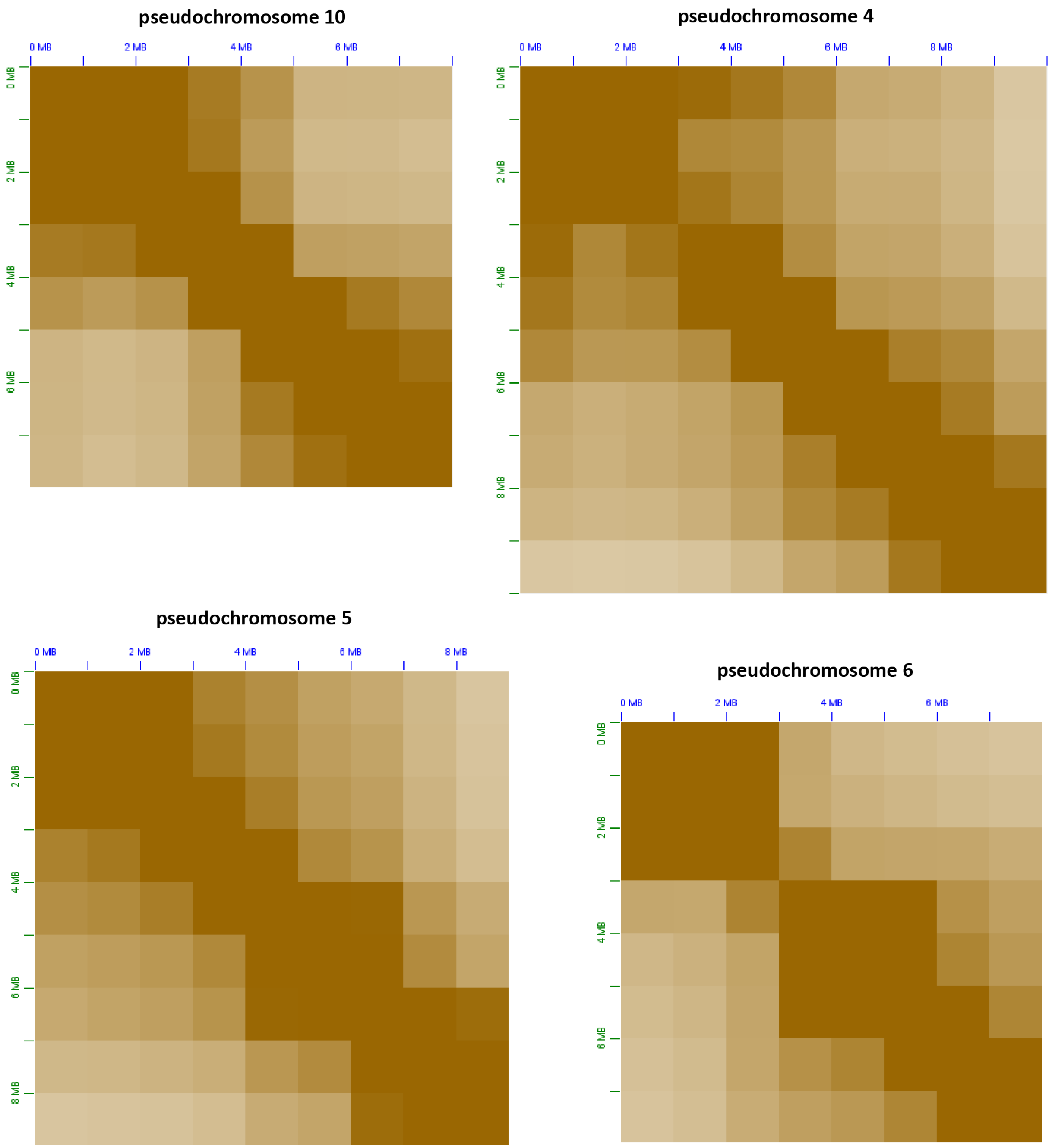


**Supplementary Figure 2: Broad-level bipartite to tripartite topological structure of dinoflagellate chromosomes**. Shown are 1Mbp-resolution KR-normalized[24] Hi-C matrices for four of the *B. minutum* pseudochromosomes.

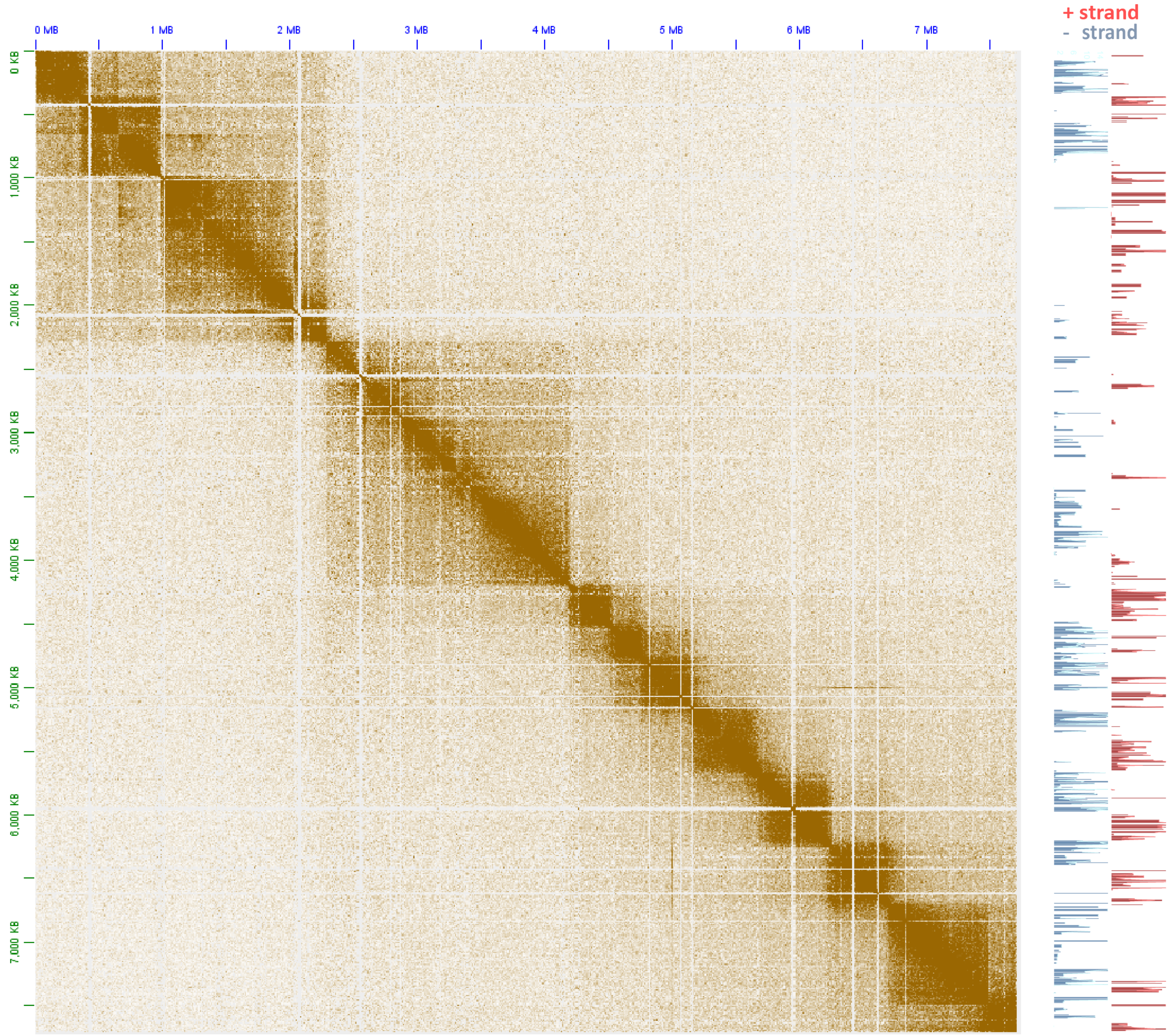


**Supplementary Figure 3: The topological domain organization of dinoflagellate chromosomes is related to polycistronic gene array orientation**. Shown is the 5kb-resolution KR-normalized Hi-C map together with strand-specific RNA expression levels for pseudochromosome 17.

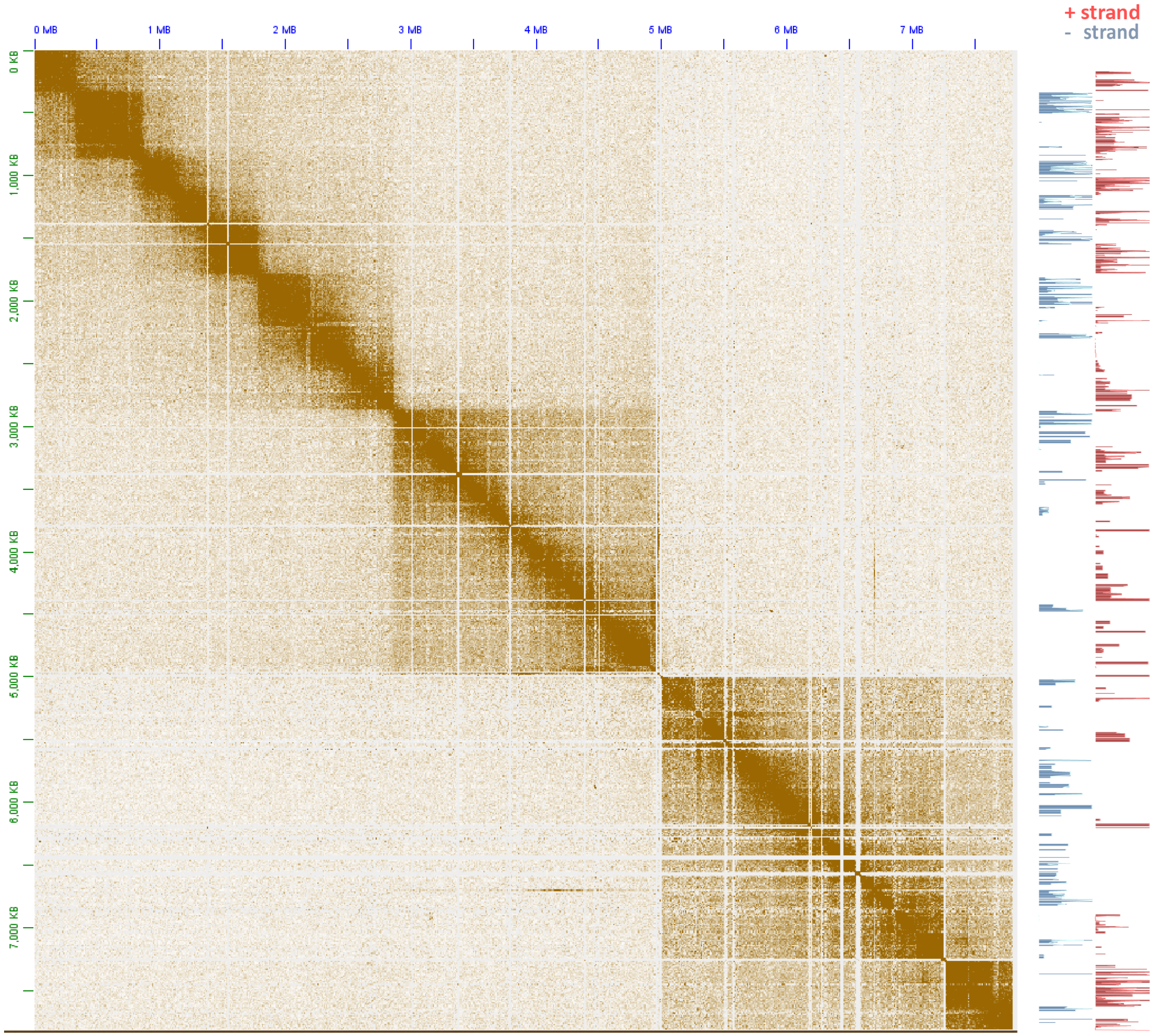


**Supplementary Figure 4: The topological domain organization of dinoflagellate chromosomes is related to polycistronic gene array orientation**. Shown is the 5kb-resolution KR-normalized Hi-C map together with strand-specific RNA expression levels for pseudochromosome 18.

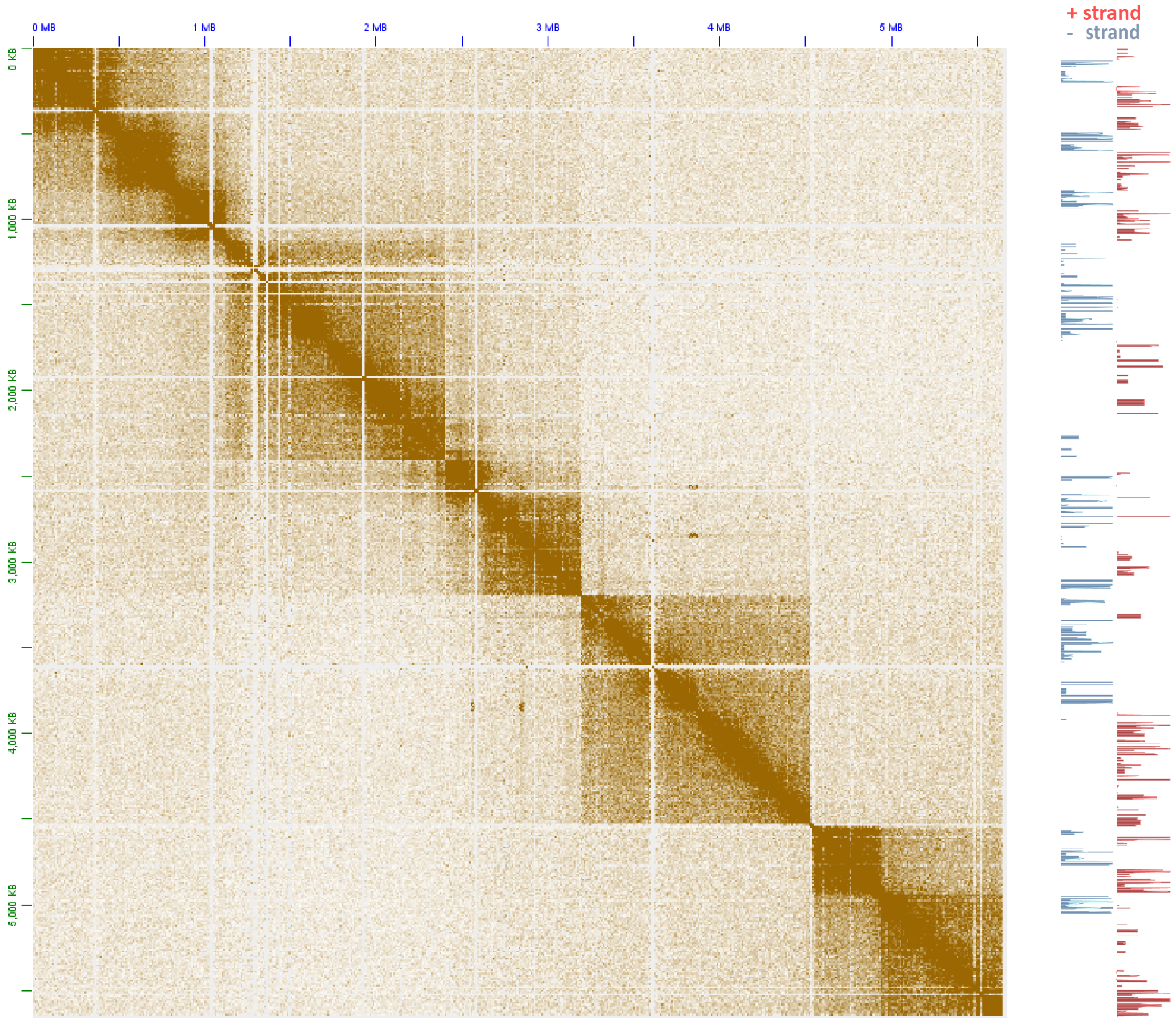


**Supplementary Figure 5: The topological domain organization of dinoflagellate chromosomes is related to polycistronic gene array orientation**. Shown is the 5kb-resolution KR-normalized Hi-C map together with strand-specific RNA expression levels for pseudochromosome 21.

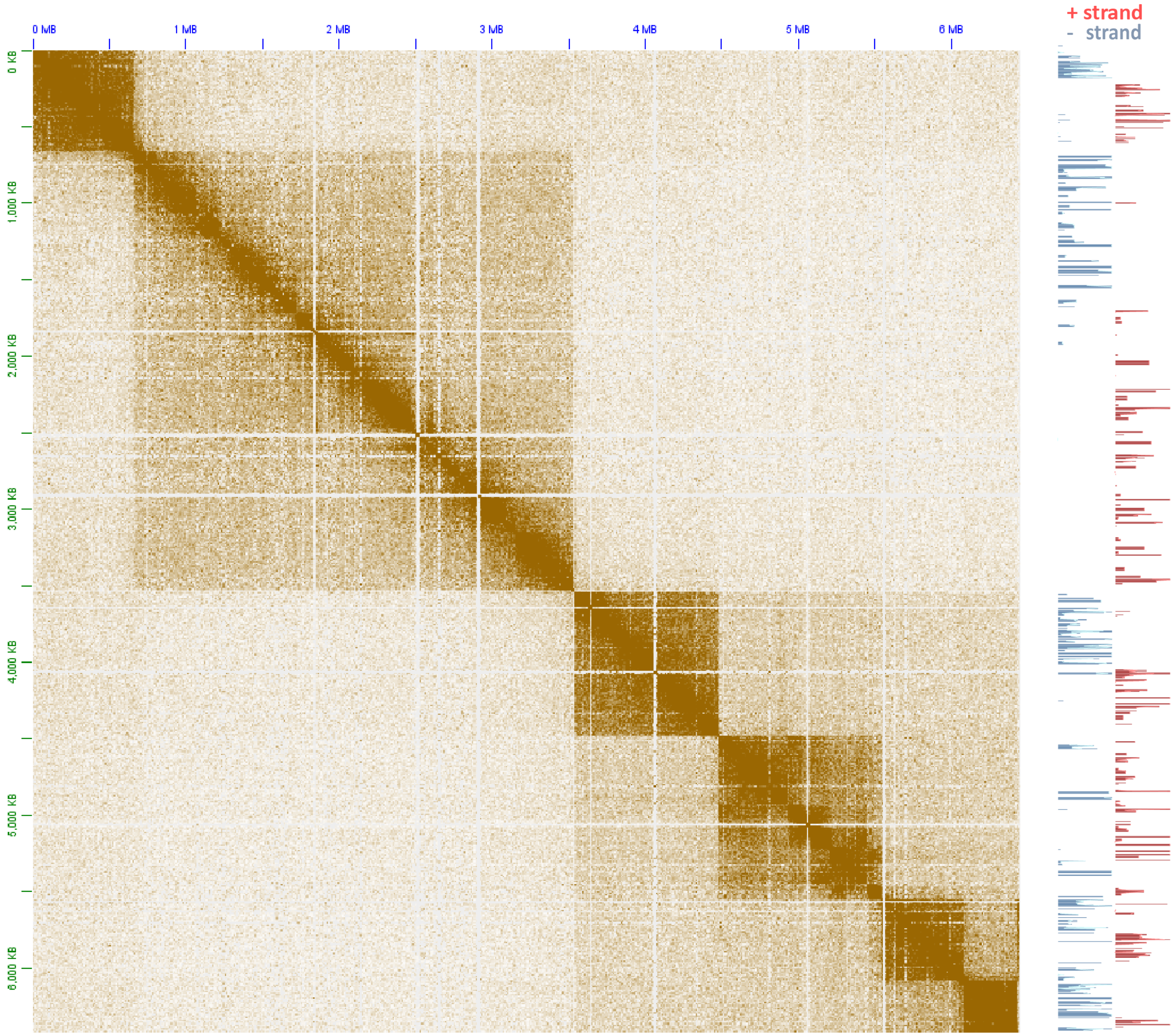


**Supplementary Figure 6: The topological domain organization of dinoflagellate chromosomes is related to polycistronic gene array orientation**. Shown is the 5kb-resolution KR-normalized Hi-C map together with strand-specific RNA expression levels for pseudochromosome 26.

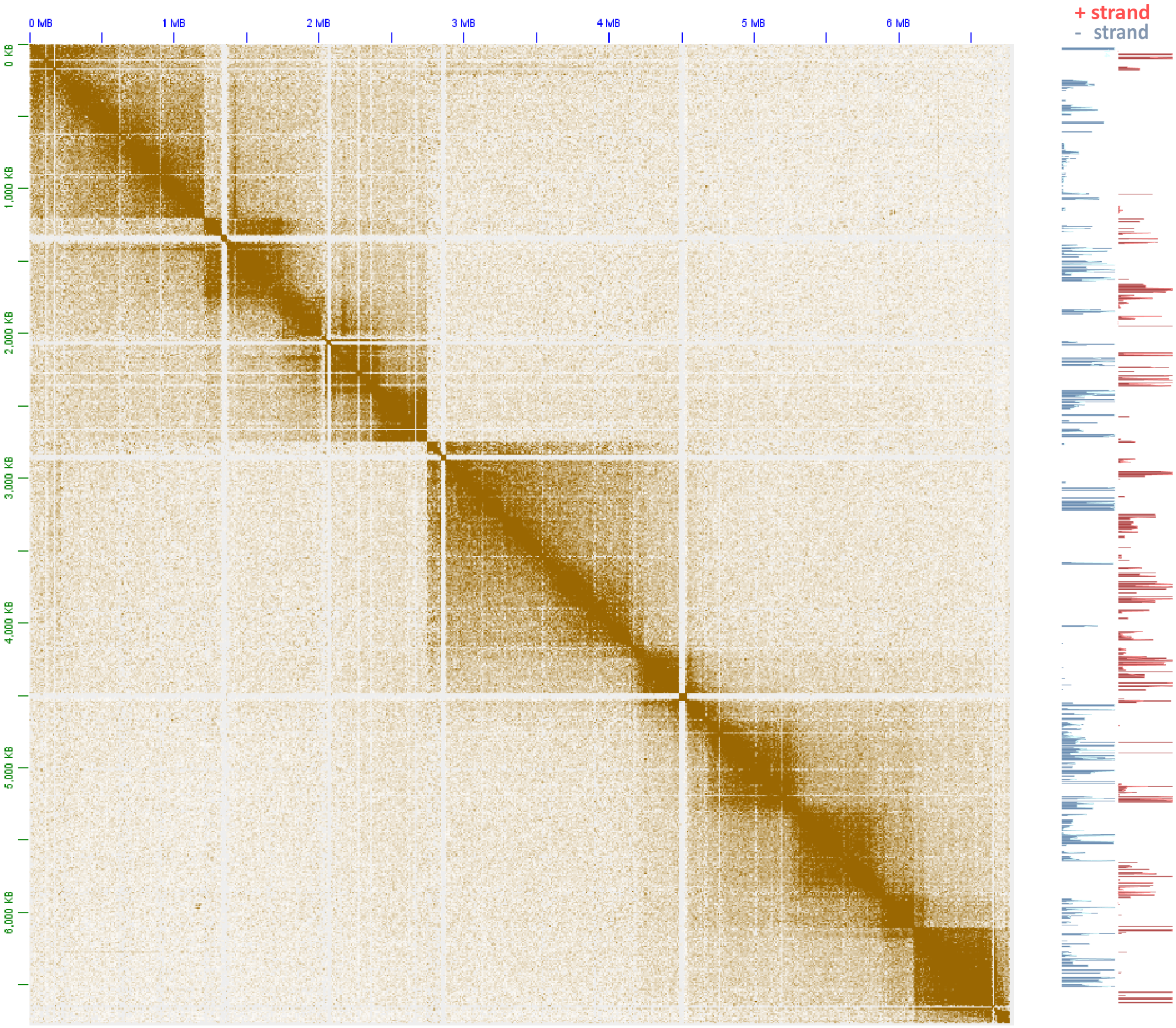


**Supplementary Figure 7: The topological domain organization of dinoflagellate chromosomes is related to polycistronic gene array orientation**. Shown is the 5kb-resolution KR-normalized Hi-C map together with strand-specific RNA expression levels for pseudochromosome 32.

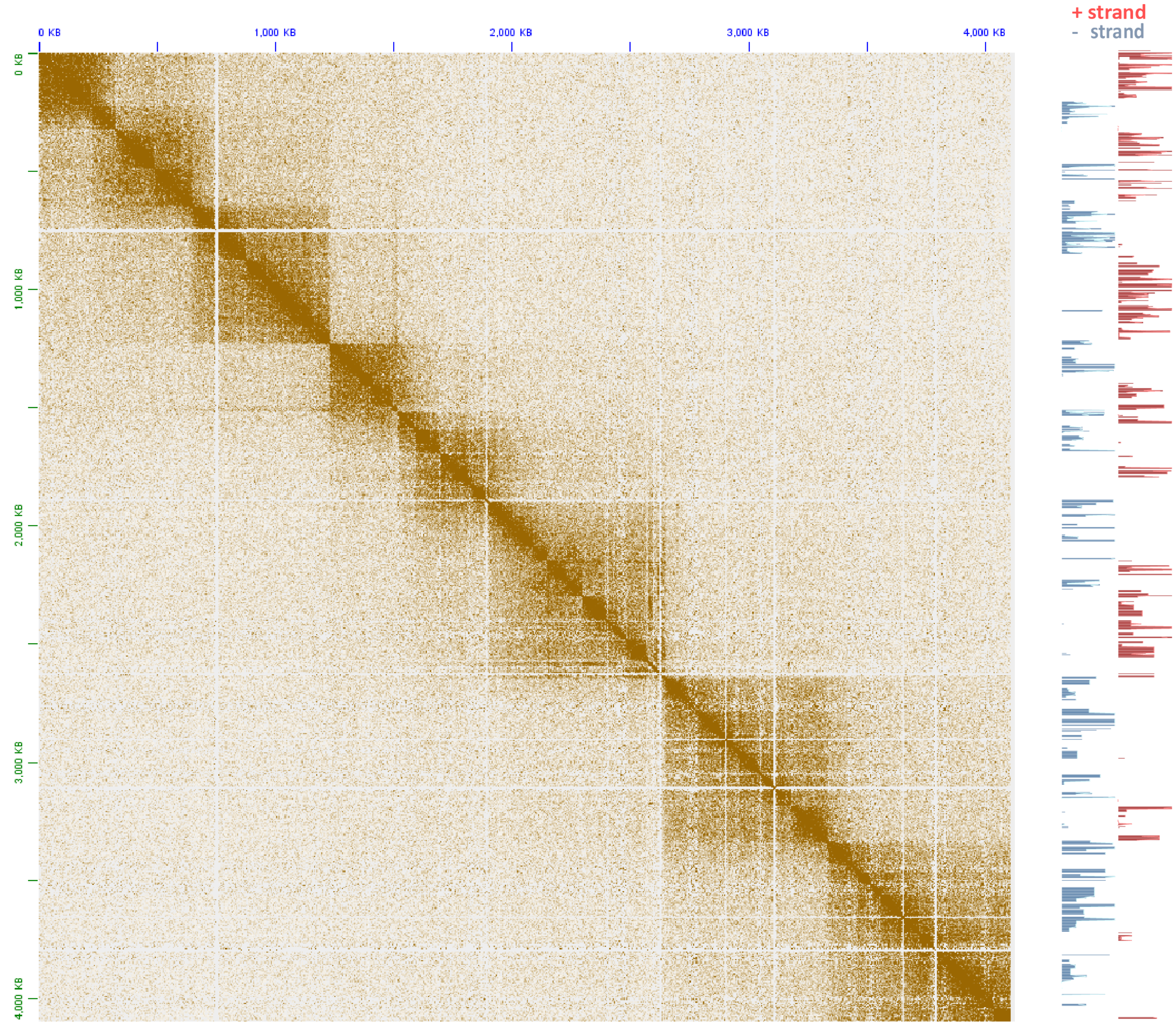


**Supplementary Figure 8: The topological domain organization of dinoflagellate chromosomes is related to polycistronic gene array orientation**. Shown is the 5kb-resolution KR-normalized Hi-C map together with strand-specific RNA expression levels for pseudochromosome 36.

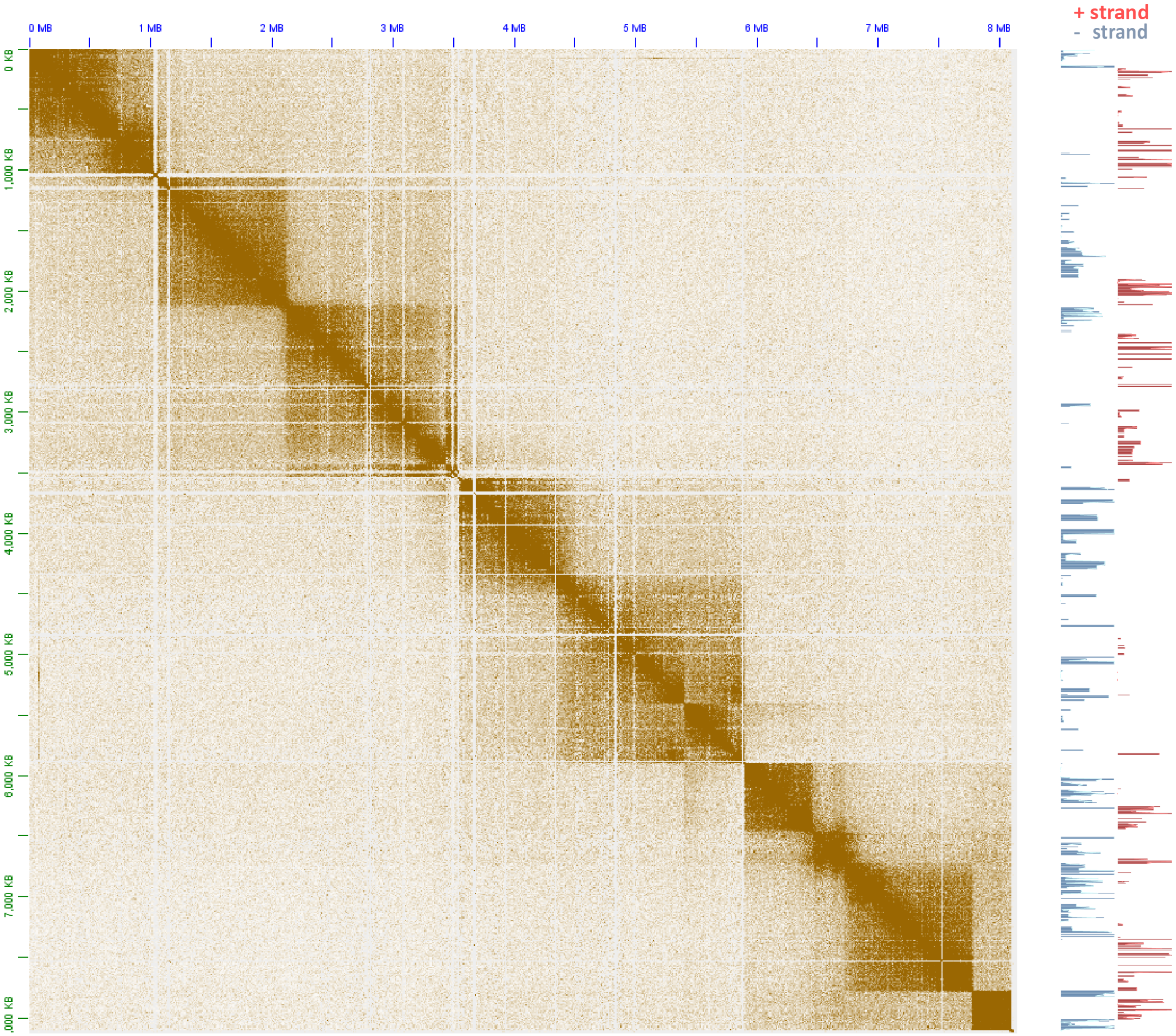


**Supplementary Figure 9: The topological domain organization of dinoflagellate chromosomes is related to polycistronic gene array orientation**. Shown is the 5kb-resolution KR-normalized Hi-C map together with strand-specific RNA expression levels for pseudochromosome 71.

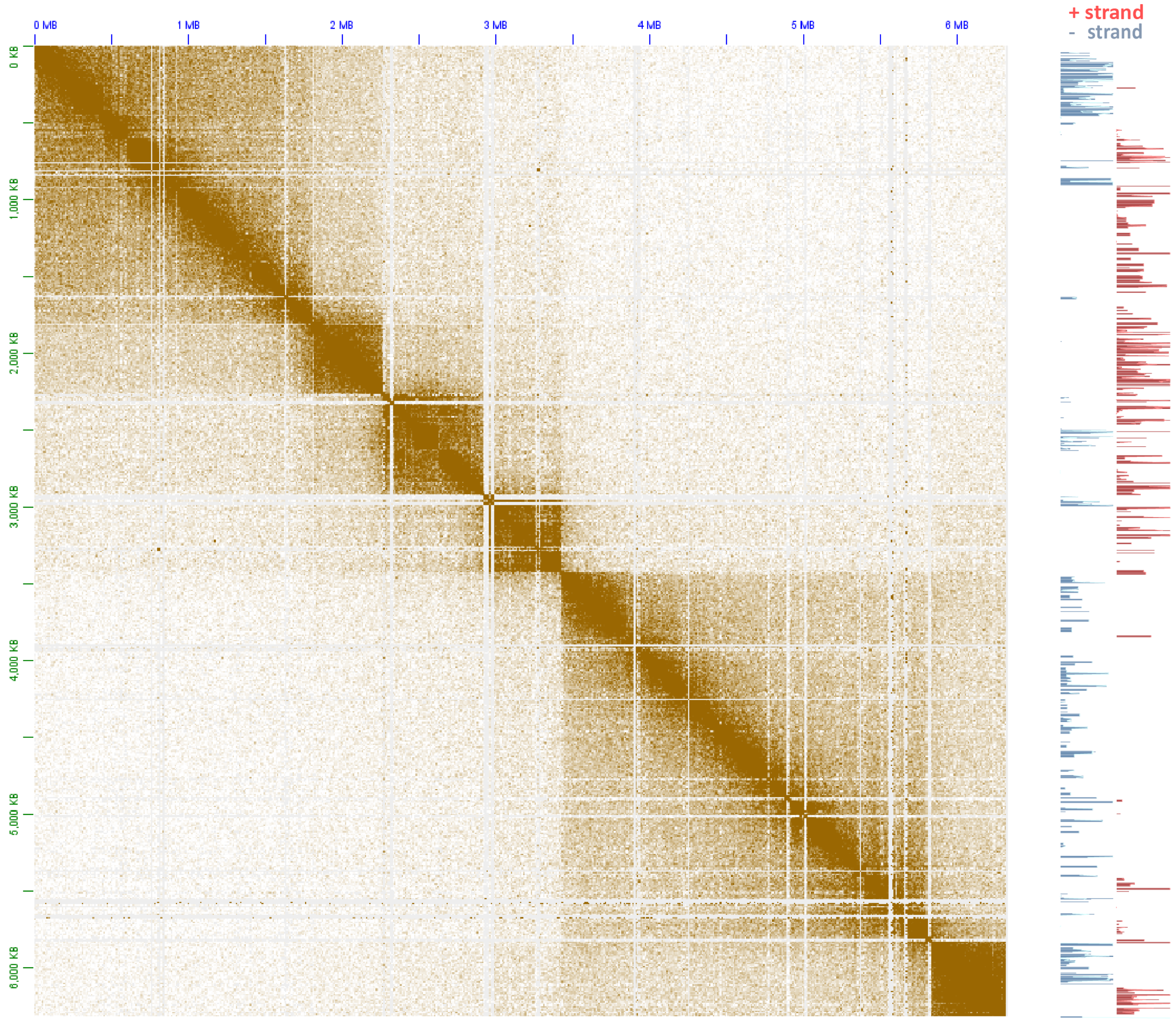


**Supplementary Figure 10: The topological domain organization of dinoflagellate chromosomes is related to polycistronic gene array orientation**. Shown is the 5kb-resolution KR-normalized Hi-C map together with strand-specific RNA expression levels for pseudochromosome 77.

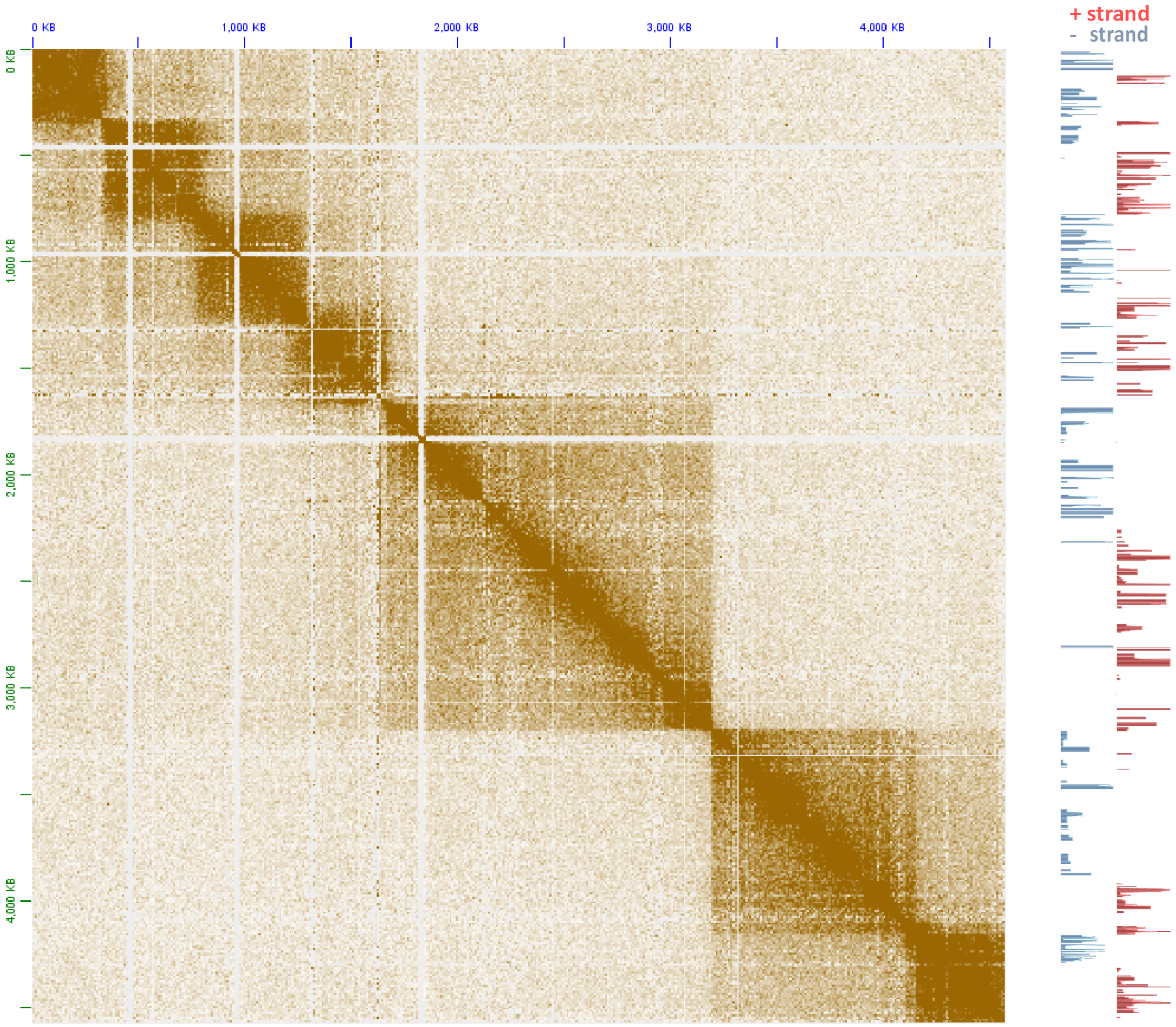


**Supplementary Figure 11: The topological domain organization of dinoflagellate chromosomes is related to polycistronic gene array orientation**. Shown is the 5kb-resolution KR-normalized Hi-C map together with strand-specific RNA expression levels for pseudochromosome 78.

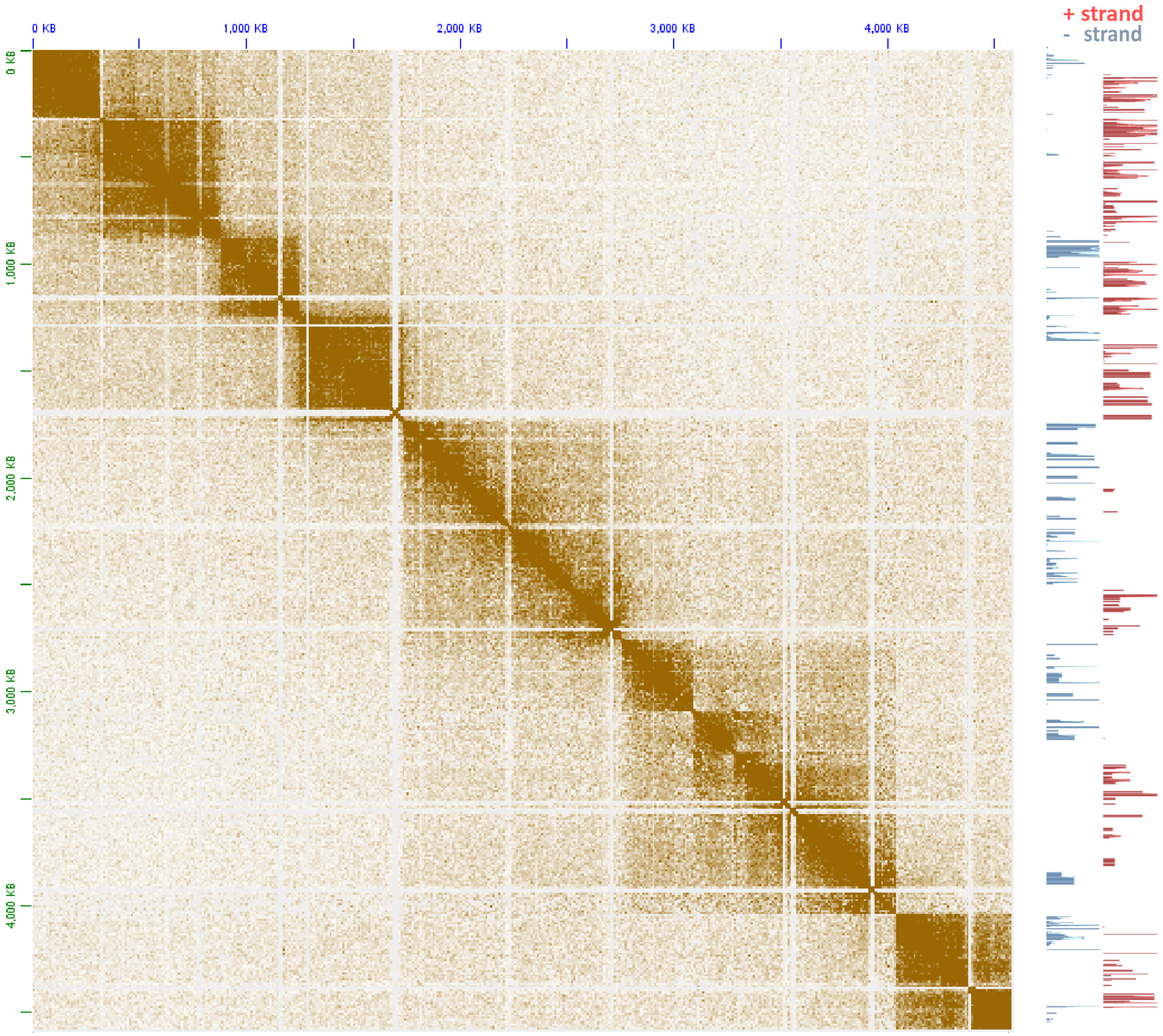


**Supplementary Figure 12: The topological domain organization of dinoflagellate chromosomes is related to polycistronic gene array orientation**. Shown is the 5kb-resolution KR-normalized Hi-C map together with strand-specific RNA expression levels for pseudochromosome 88.

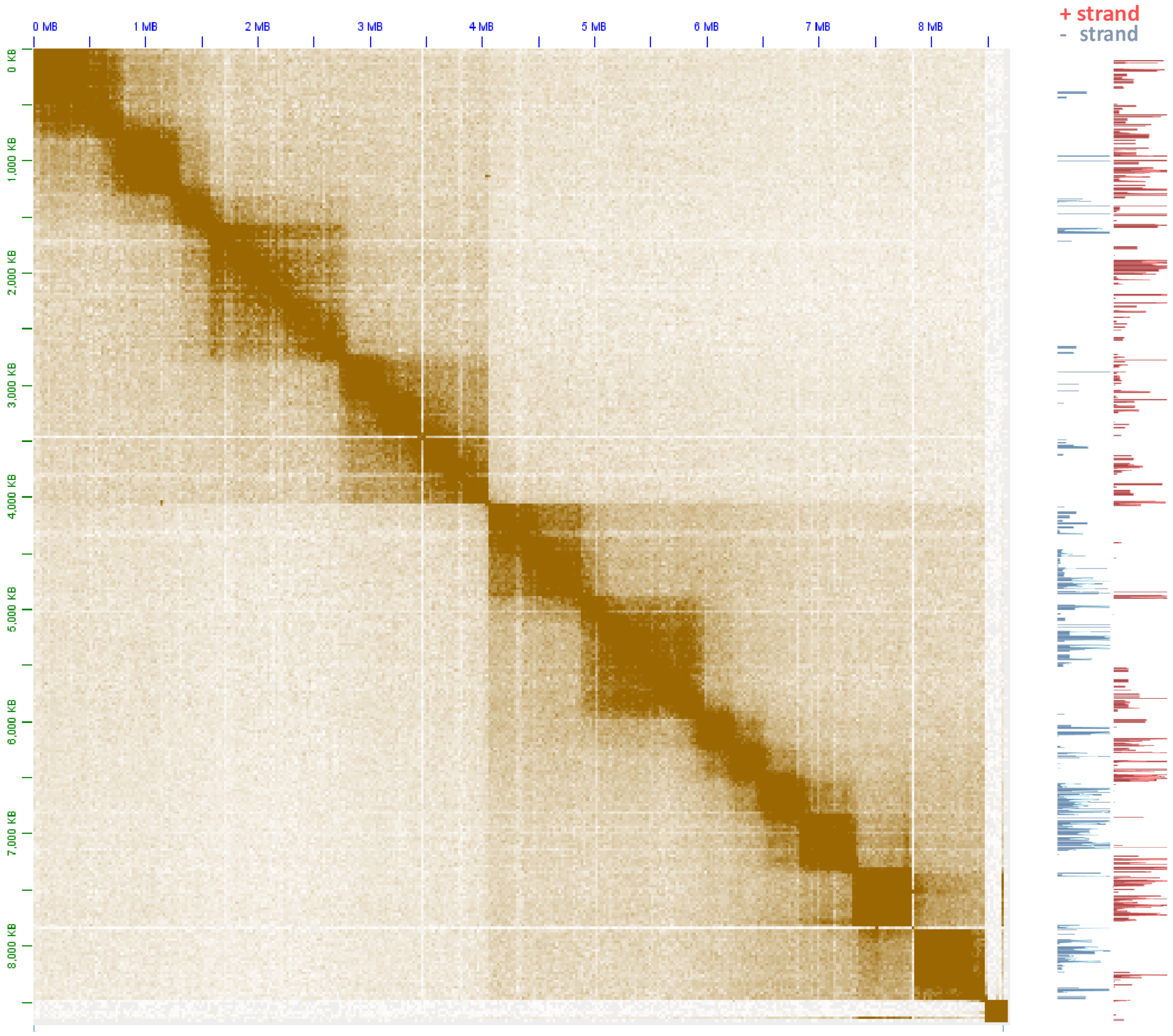


**Supplementary Figure 13: The topological domain organization of dinoflagellate chromosomes is related to polycistronic gene array orientation**. Shown is the 5kb-resolution KR-normalized Hi-C map together with strand-specific RNA expression levels for pseudochromosome 89.

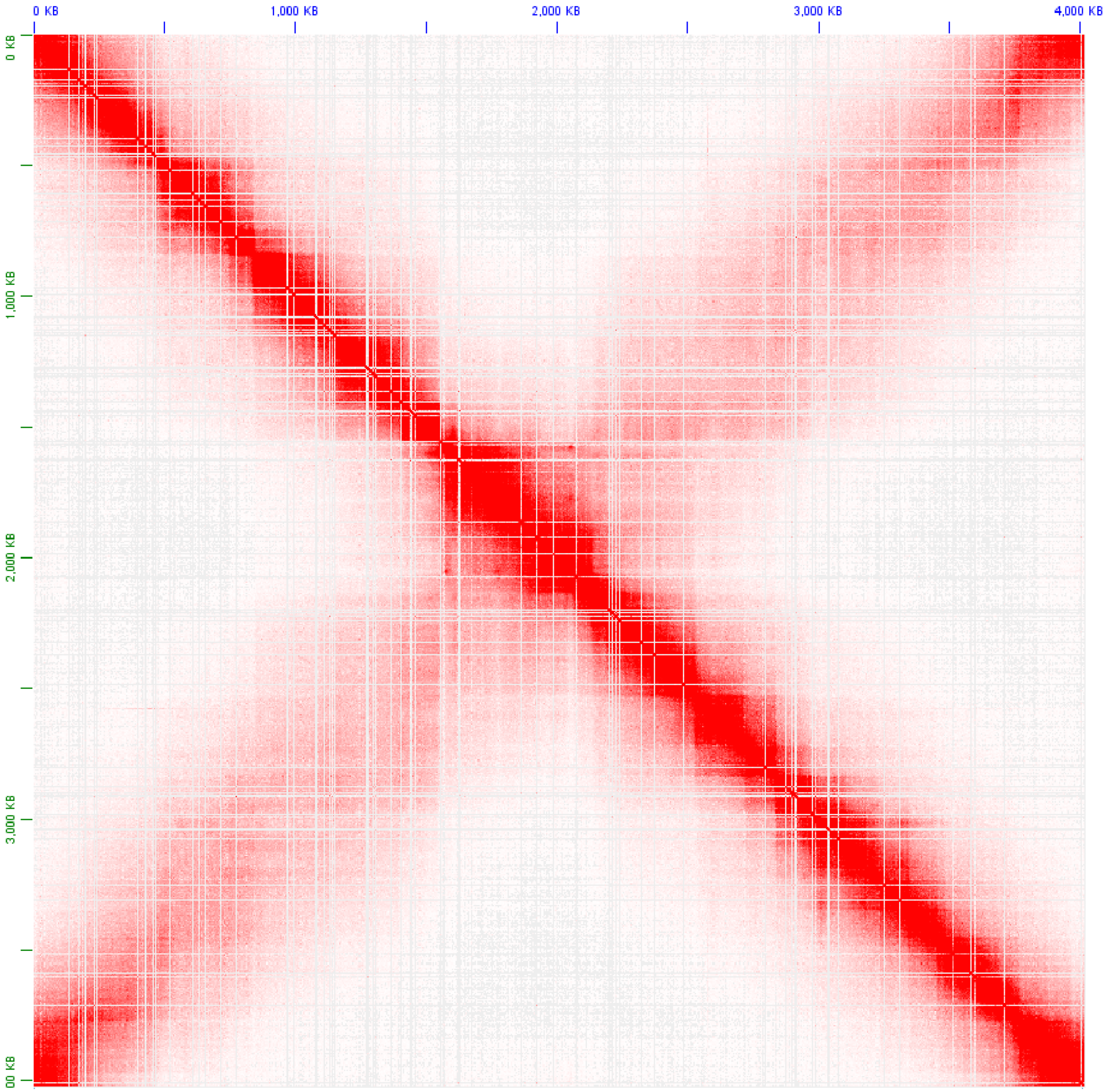


**Supplementary Figure 14: Topological structure of the *Caulobacter crescentus* CB15 genome**. Shown is the KR-normalized 5-kb resolution maps for the whole *Caulobacter* chromosome (GEO accession GSM1120448).

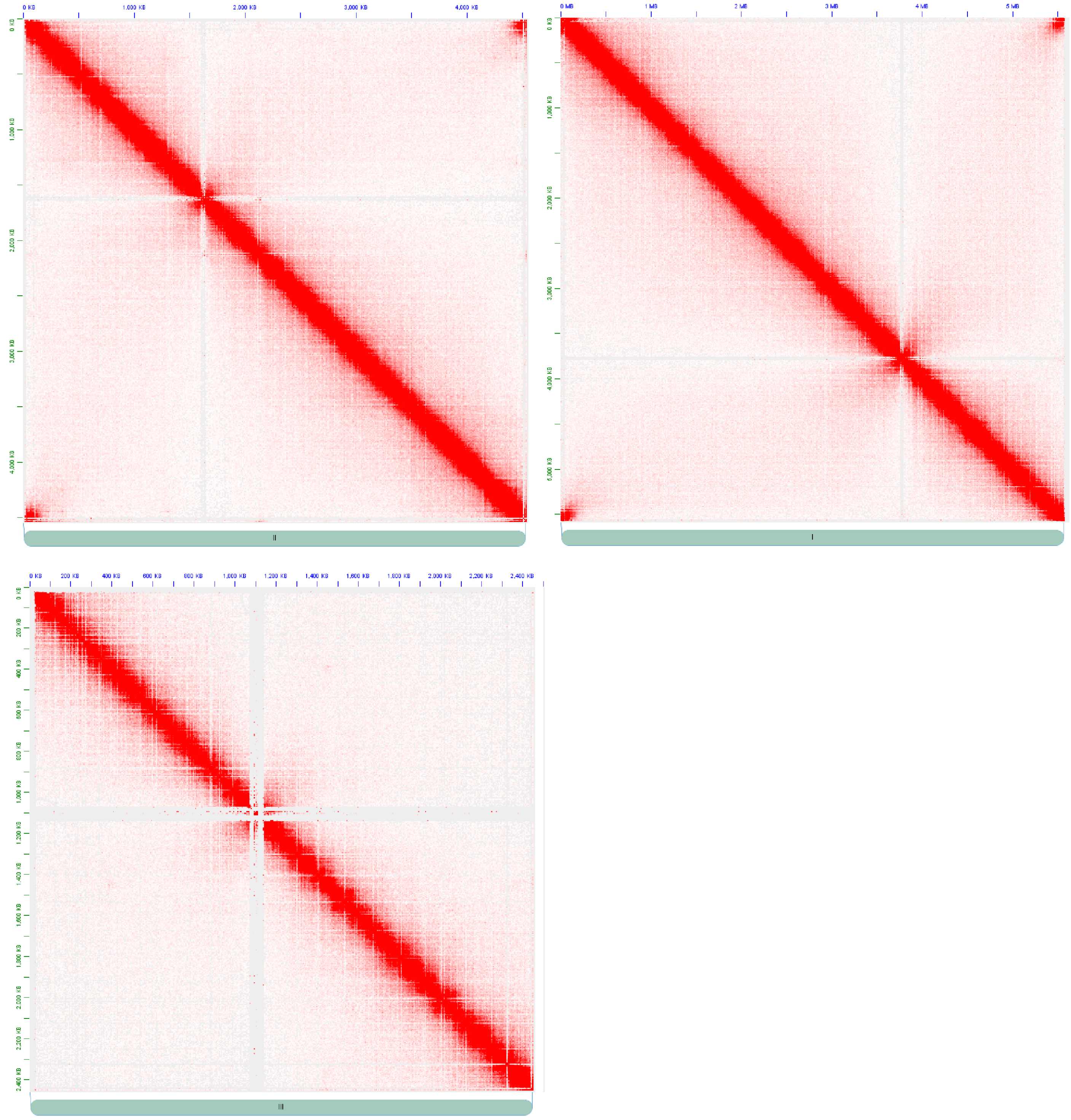


**Supplementary Figure 15: Topological structure of the *Schizosaccharomyces pombe* genome**. Shown are the KR-normalized 5-kb resolution maps for all three *S. pombe* chromosome (GEO accession GSM1379427).

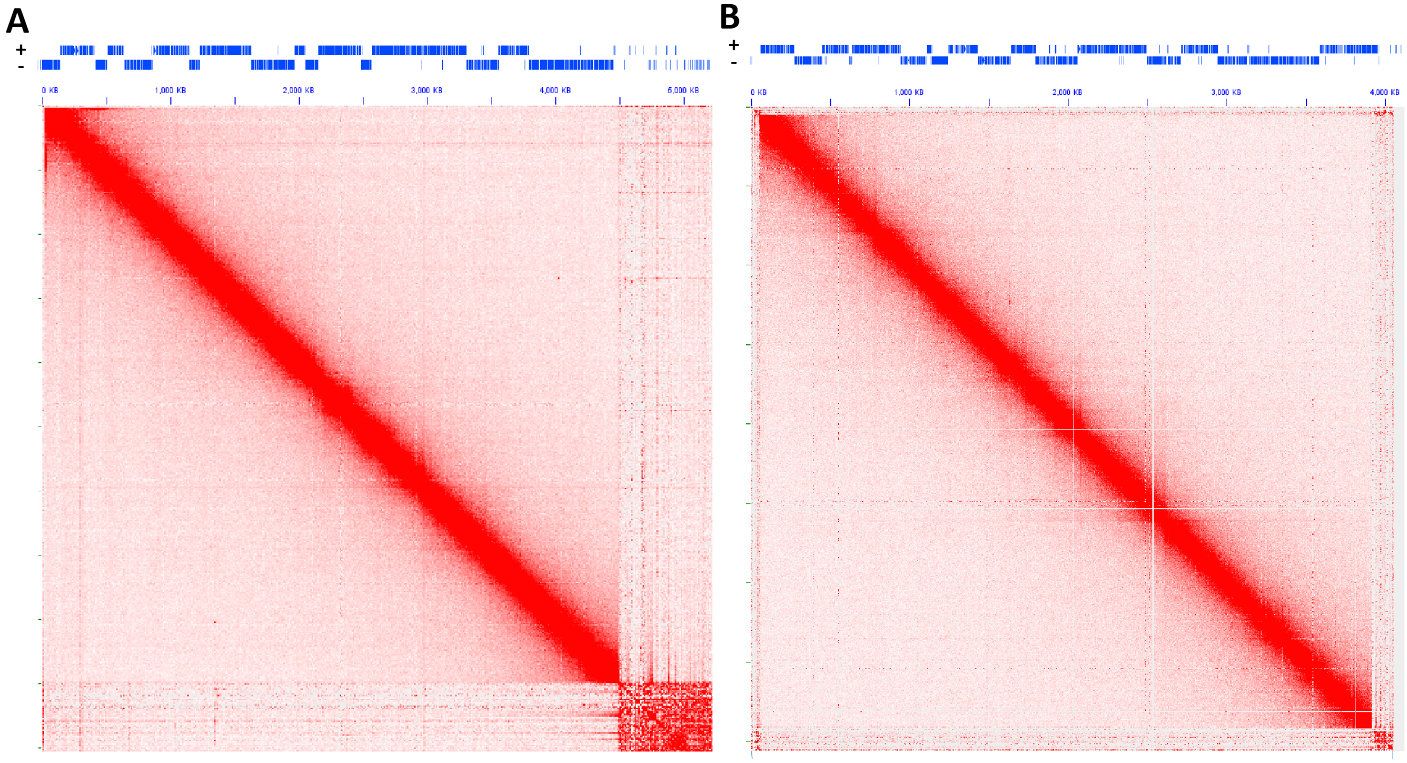


**Supplementary Figure 16: No topological domains associated with gene arrays are observed in the kinetoplastid *Trypanosoma brucei*.** Shown are KR-normalized 10-kb resolution maps for chr11 (A) and chr10 (B) for GEO accession GSM3346690.

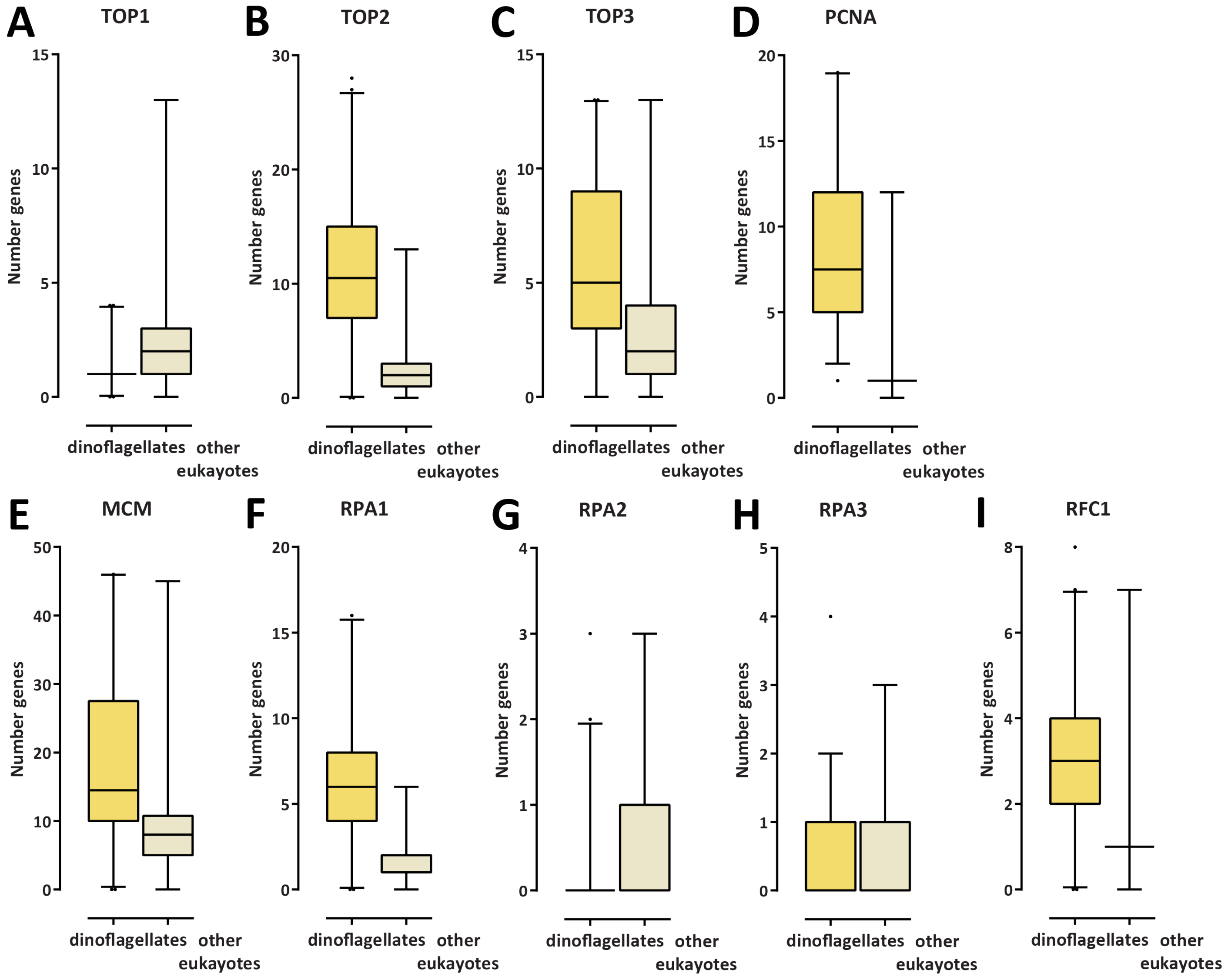


**Supplementary Figure 17: Expansion of the Type II and II topoisomerase gene repertoire as well as of certain other replication-related proteins in dinoflagellates**. Shown are the number of genes annotated in MMETSP transcriptome assemblies of dinoflagellates and other eukaryotes. (A) Number of Type I topoisomerase genes; (B) Number of Type II topoisomerase genes; (C) Number of Type III topoisomerase genes; (D) Number of PCNA genes; (E) Number of MCM genes; (F) Number of RPA1 genes; (G) Number of RPA2 genes; (H) Number of RPA3 genes; (I) Number of RFC1 genes.

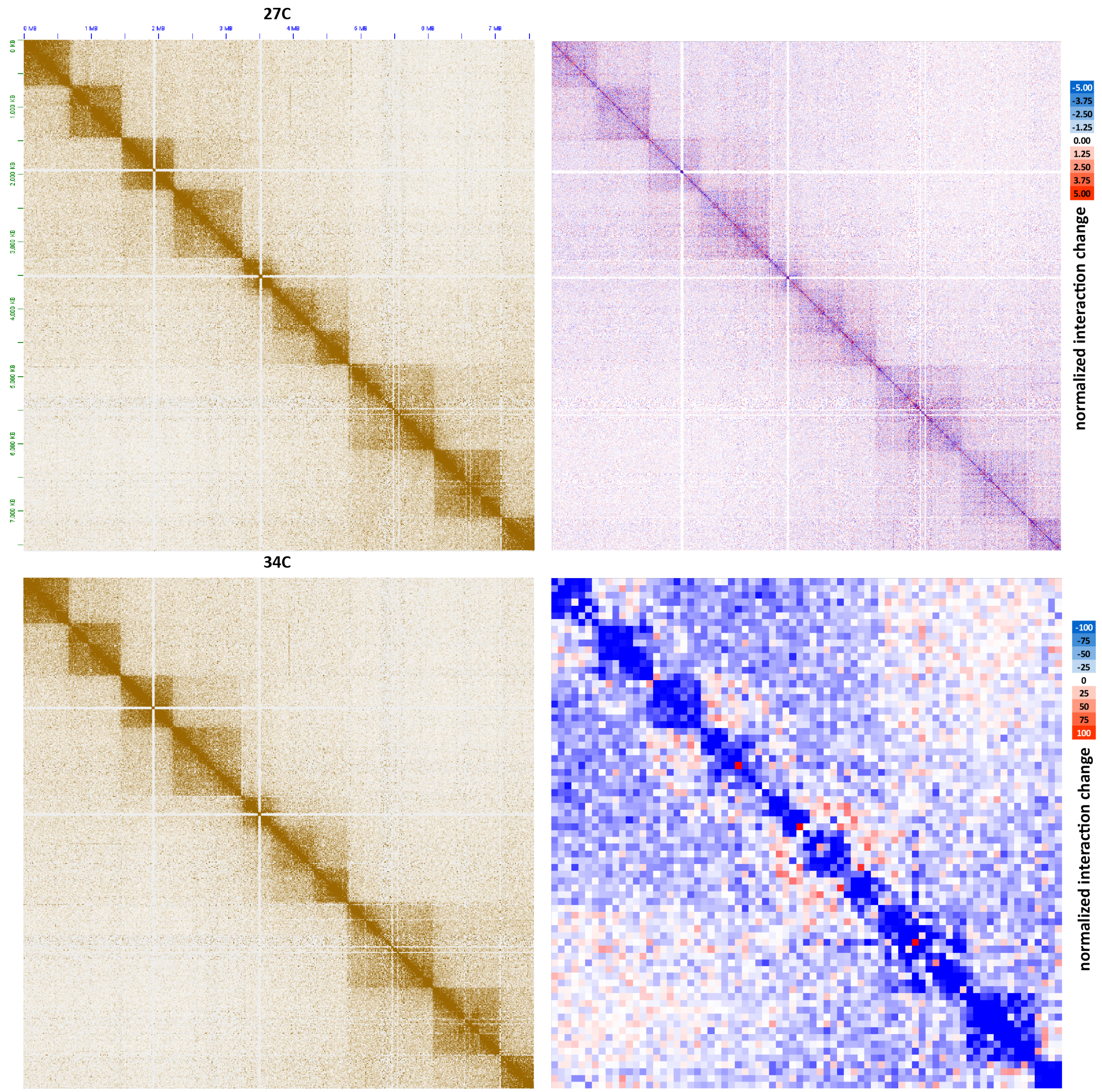


**Supplementary Figure 18: Moderate decompaction of dinoTADs upon exposure to elevated temperatures**. Shown is pseudochromosome 10 (KR-normalized) and the difference between the KR-normalized Hi-C maps generated from *B. minutum* grown at 34 °C and at 27 °C at 100-kb resolution (lower right) and 5-kb resolution (upper right).

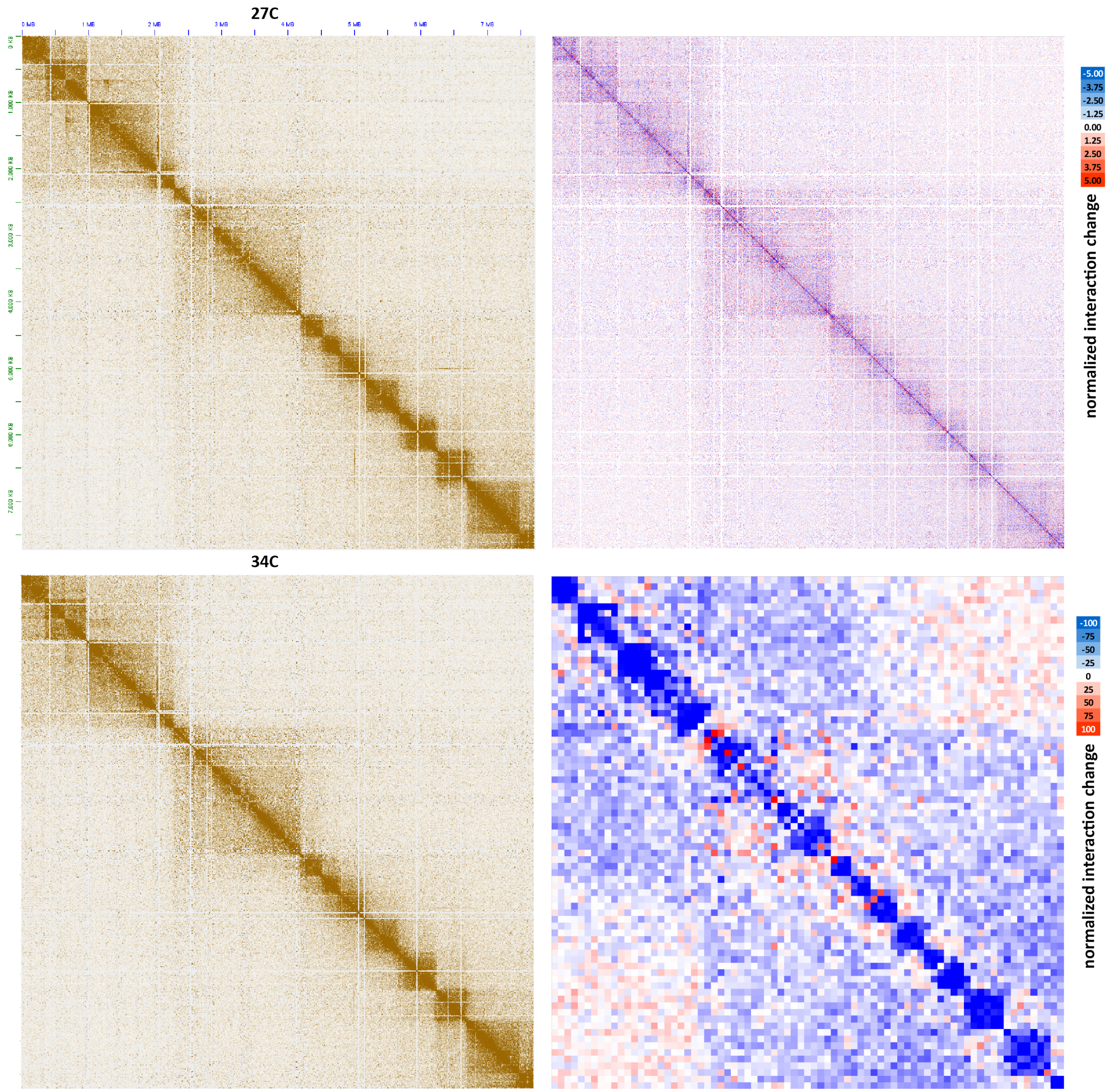


**Supplementary Figure 19: Moderate decompaction of dinoTADs upon exposure to elevated temperatures.** Shown is pseudochromosome 17 (KR-normalized) and the difference between the KR-normalized Hi-C maps generated from *B. minutum* grown at 34 °C and at 27 °C at 100-kb resolution (lower right) and 5-kb resolution (upper right).

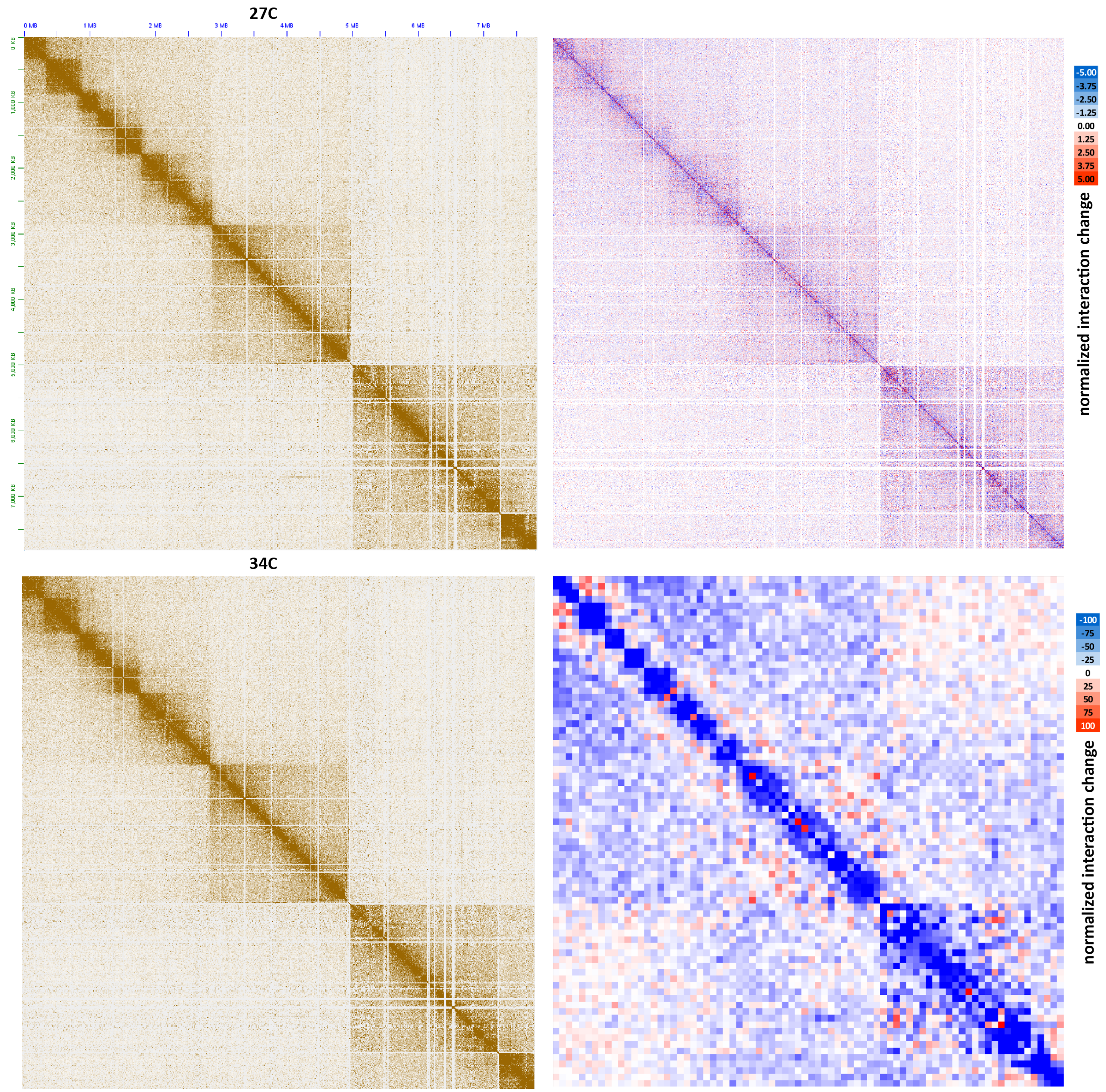


**Supplementary Figure 20: Moderate decompaction of dinoTADs upon exposure to elevated temperatures**. Shown is pseudochromosome 18 (KR-normalized) and the difference between the KR-normalized Hi-C maps generated from *B. minutum* grown at 34 °C and at 27 °C at 100-kb resolution (lower right) and 5-kb resolution (upper right).

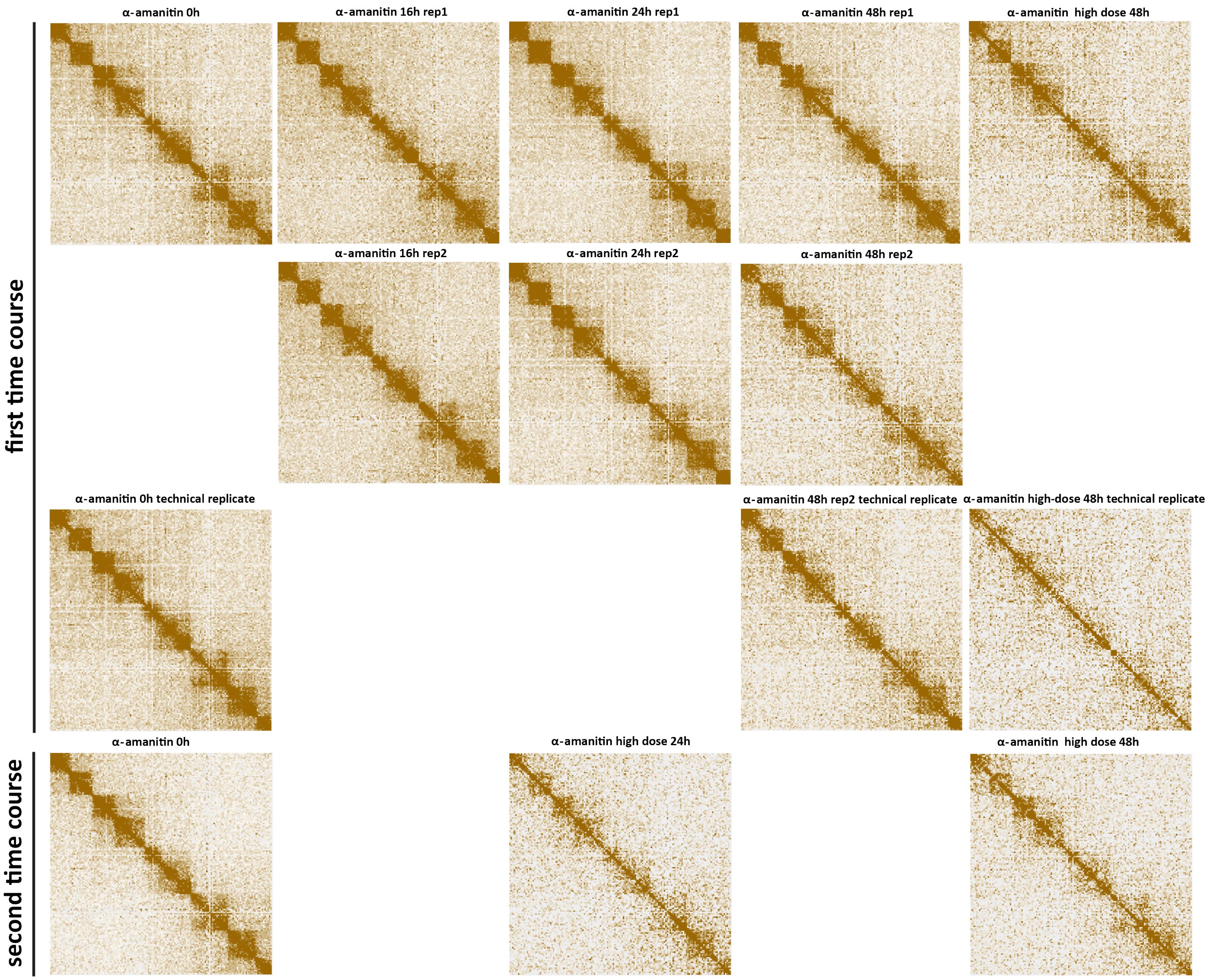


**Supplementary Figure 21: Decompaction of dinoTADs upon transcriptional inhibition using $\alpha$-amanitin.** Shown is pseudochromosome 10. Two time courses were carried out following the outline presented in Figure 2B.

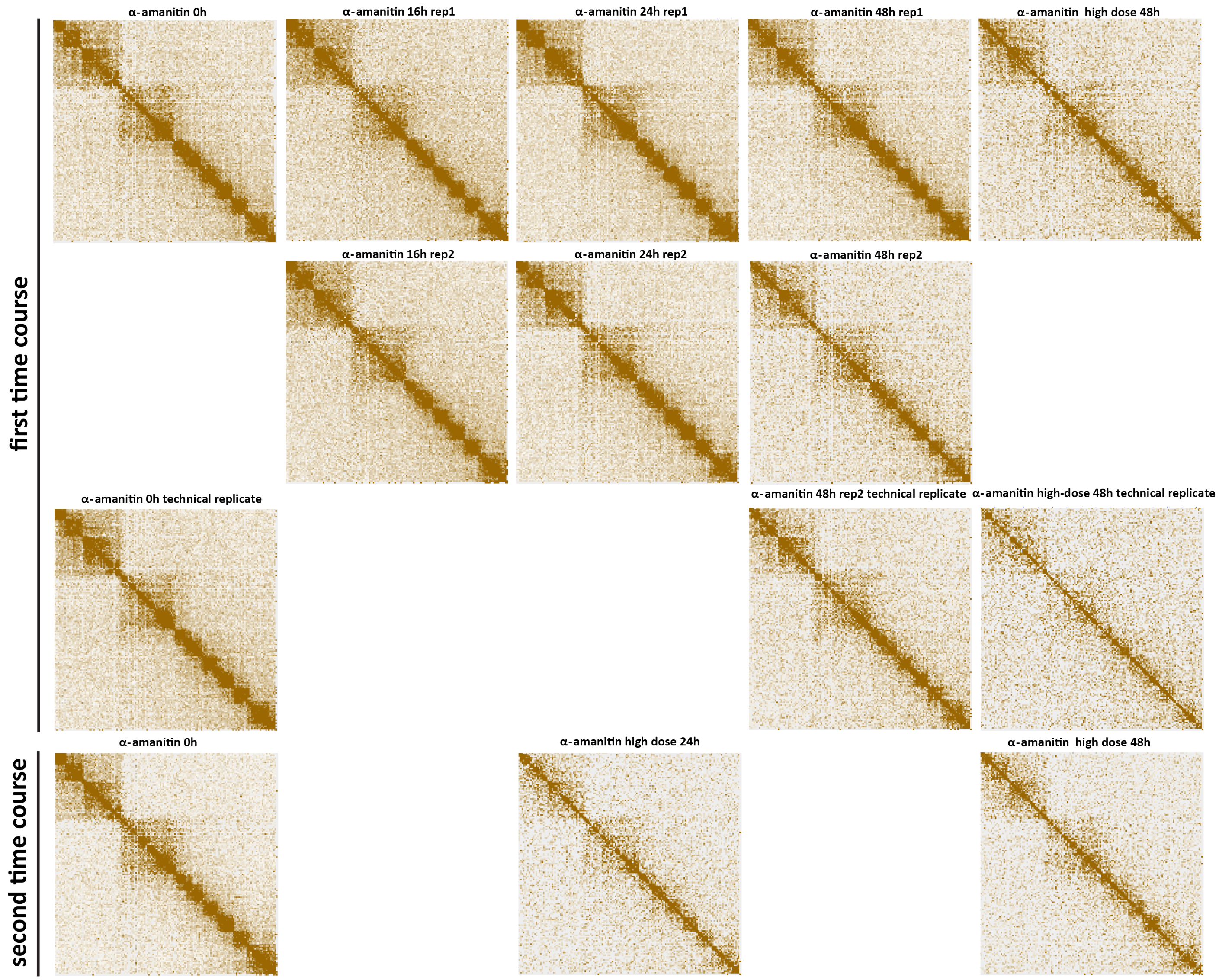


**Supplementary Figure 22: Decompaction of dinoTADs upon transcriptional inhibition using $\alpha$-amanitin.** Shown is pseudochromosome 17. Two time courses were carried out following the outline presented in Figure 2B.

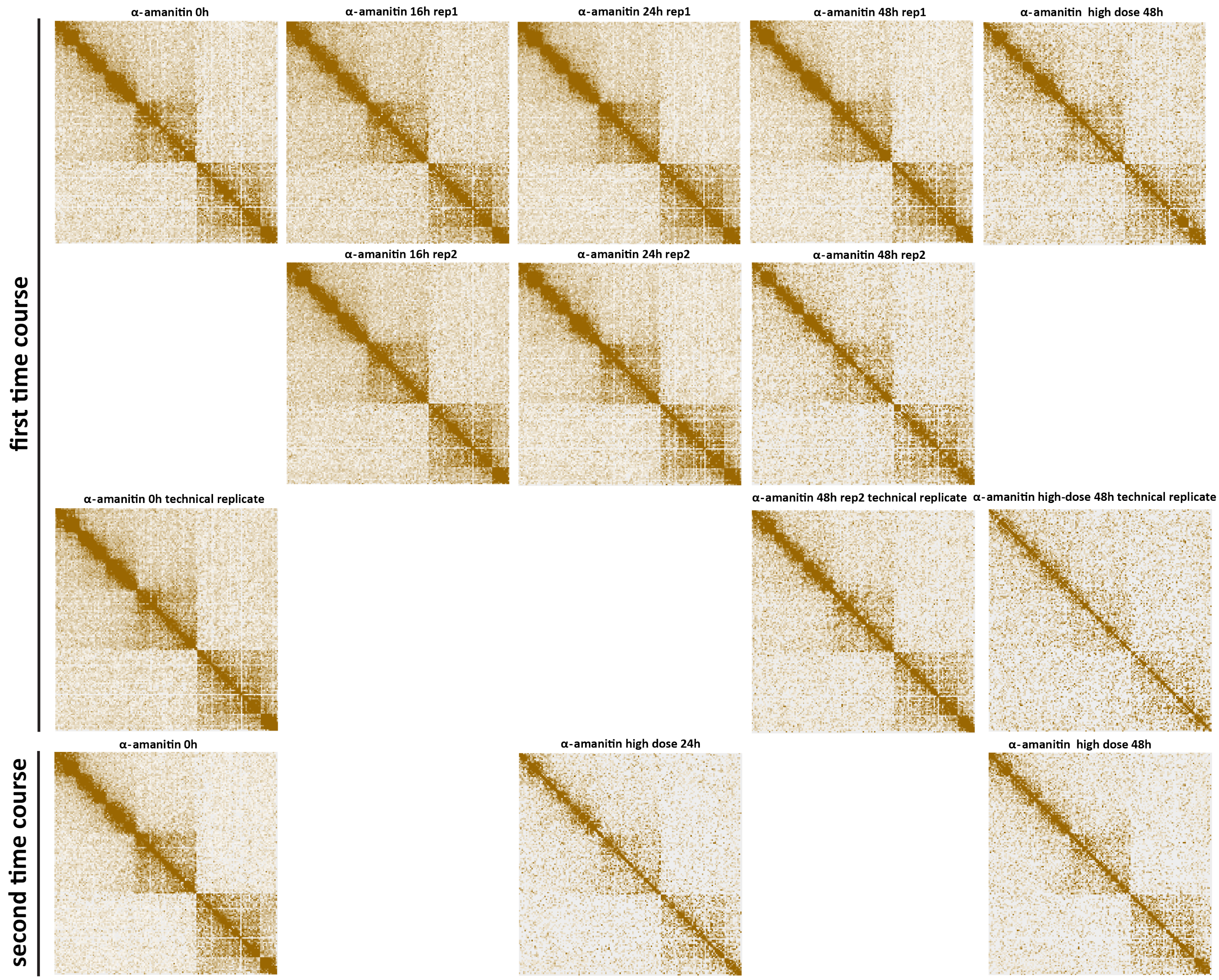


**Supplementary Figure 23: Decompaction of dinoTADs upon transcriptional inhibition using $\alpha$-amanitin.** Shown is pseudochromosome 18. Two time courses were carried out following the outline presented in Figure 2B.

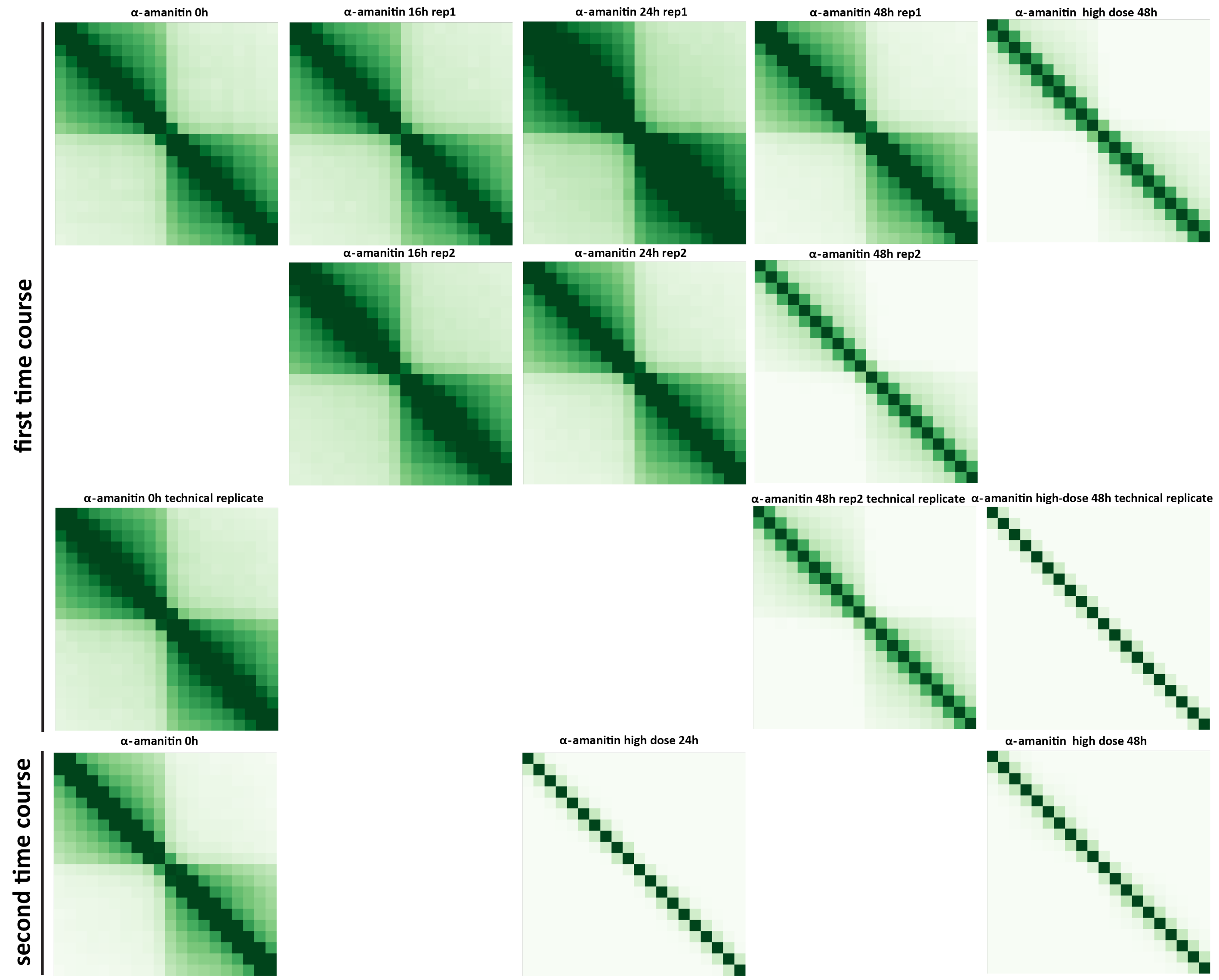


**Supplementary Figure 24: Decompaction of dinoTADs upon transcriptional inhibition using $\alpha$-amanitin.** Shown are 50-kb resolution metaplots centered on dinoTAD domain boundaries. Two time courses were carried out following the outline presented in Figure 2B.

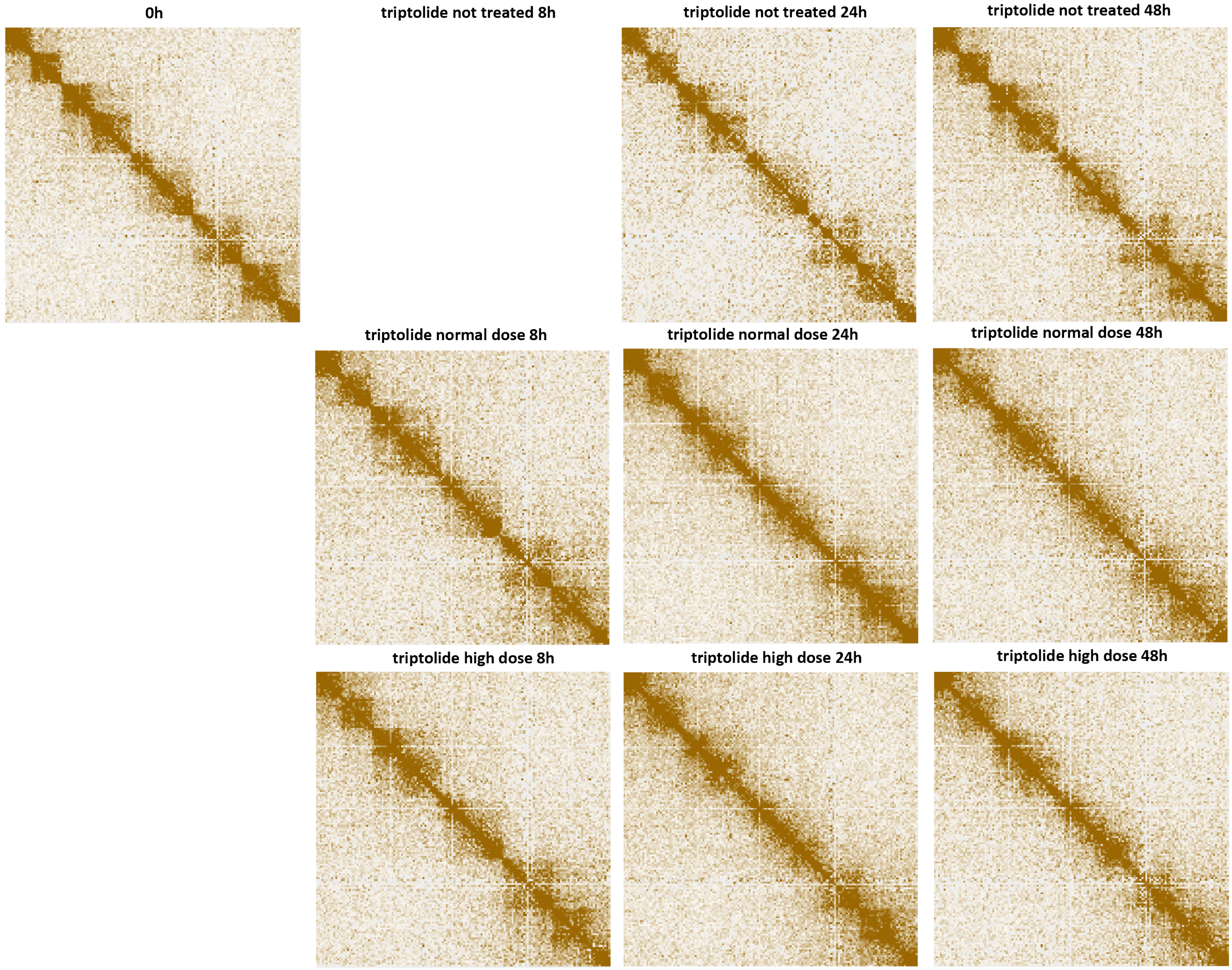


**Supplementary Figure 25: Blurring of dinoTAD boundaries upon transcriptional inhibition using triptolide**. Shown is pseudochromosome 10. The triptolide time course was carried out following the outline presented in Figure 2B.

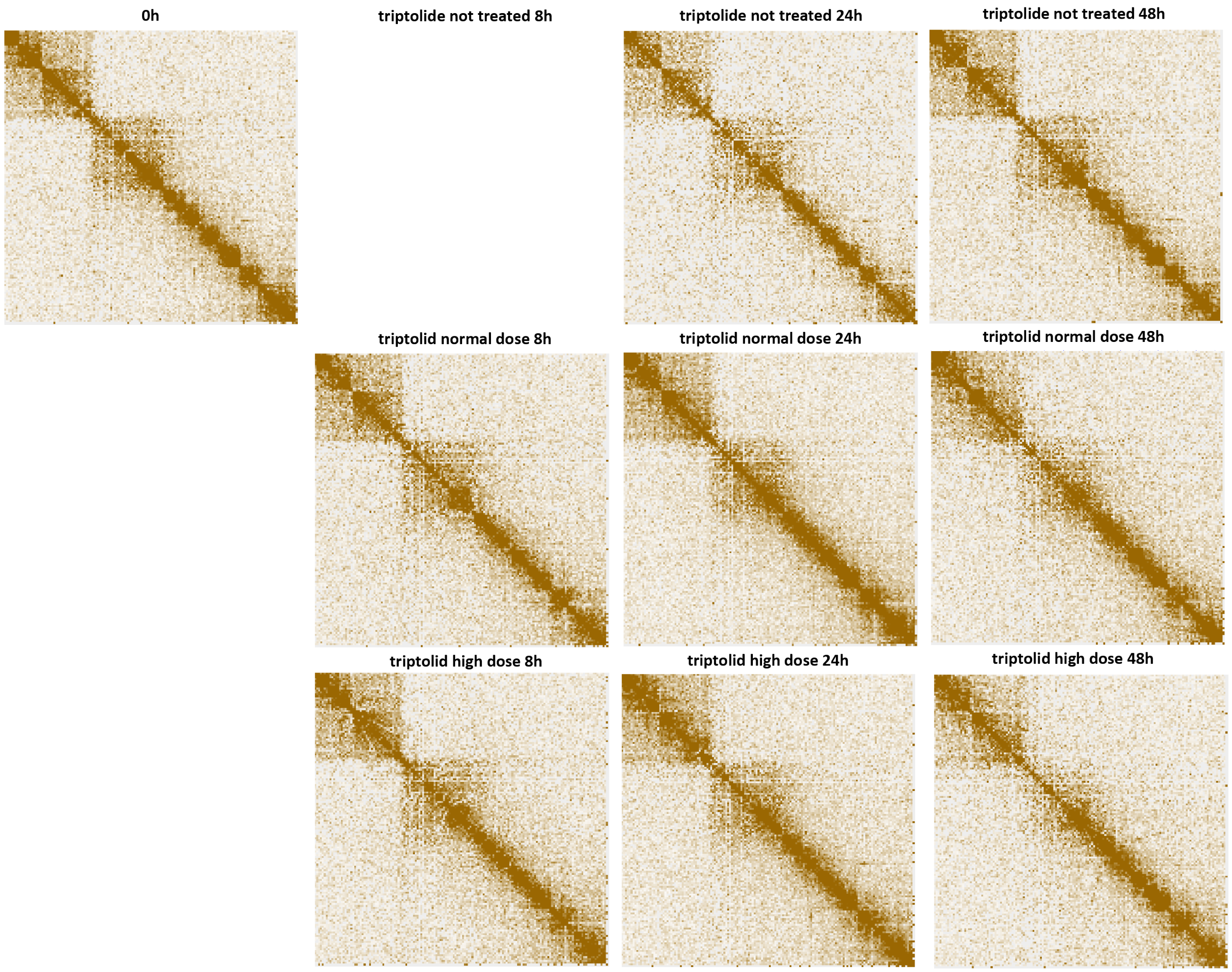


**Supplementary Figure 26: Blurring of dinoTAD boundaries upon transcriptional inhibition using triptolide**. Shown is pseudochromosome 17. The triptolide time course was carried out following the outline presented in Figure 2B.

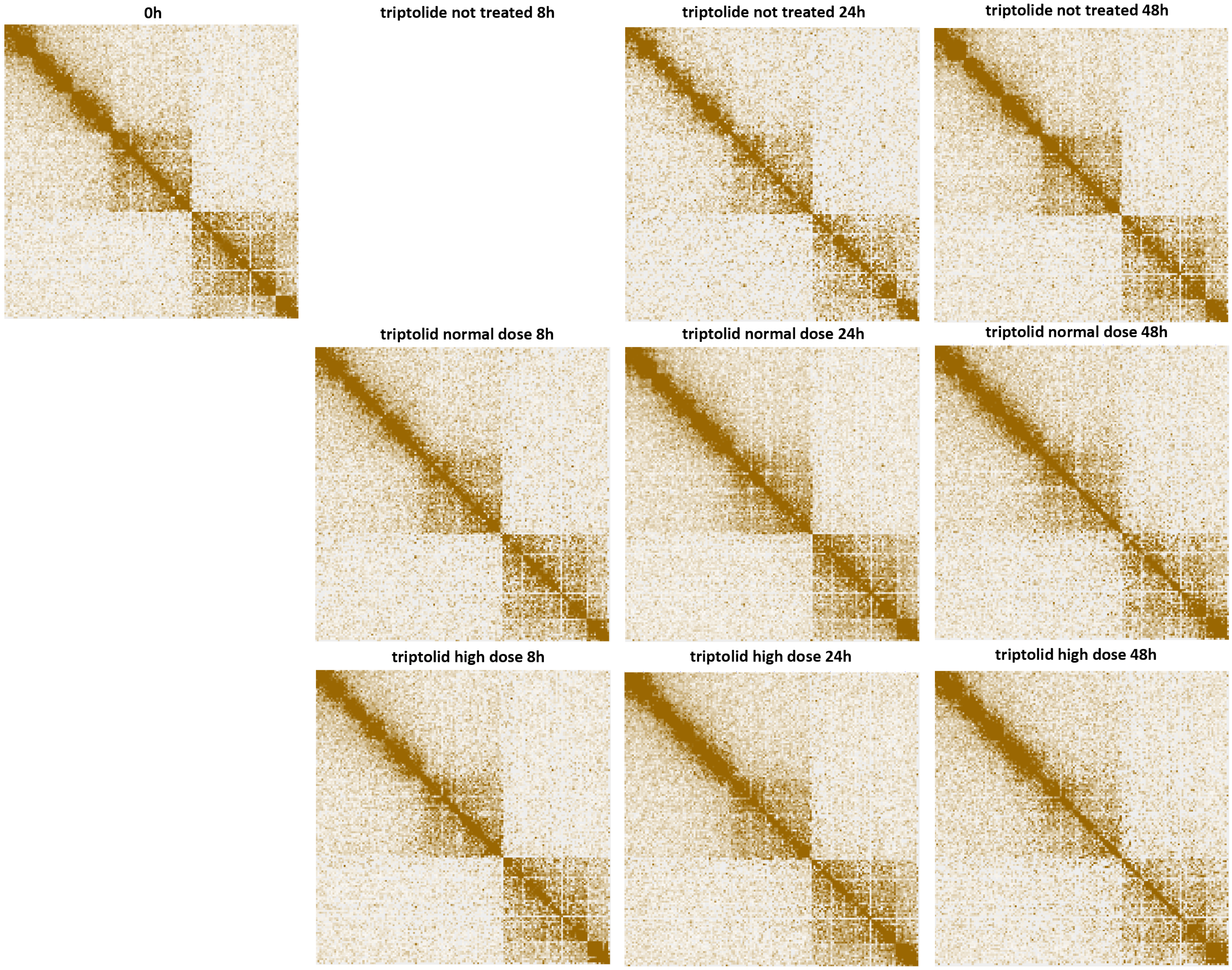


**Supplementary Figure 27: Blurring of dinoTAD boundaries upon transcriptional inhibition using triptolide**. Shown is pseudochromosome 18. The triptolide time course was carried out following the outline presented in Figure 2B.

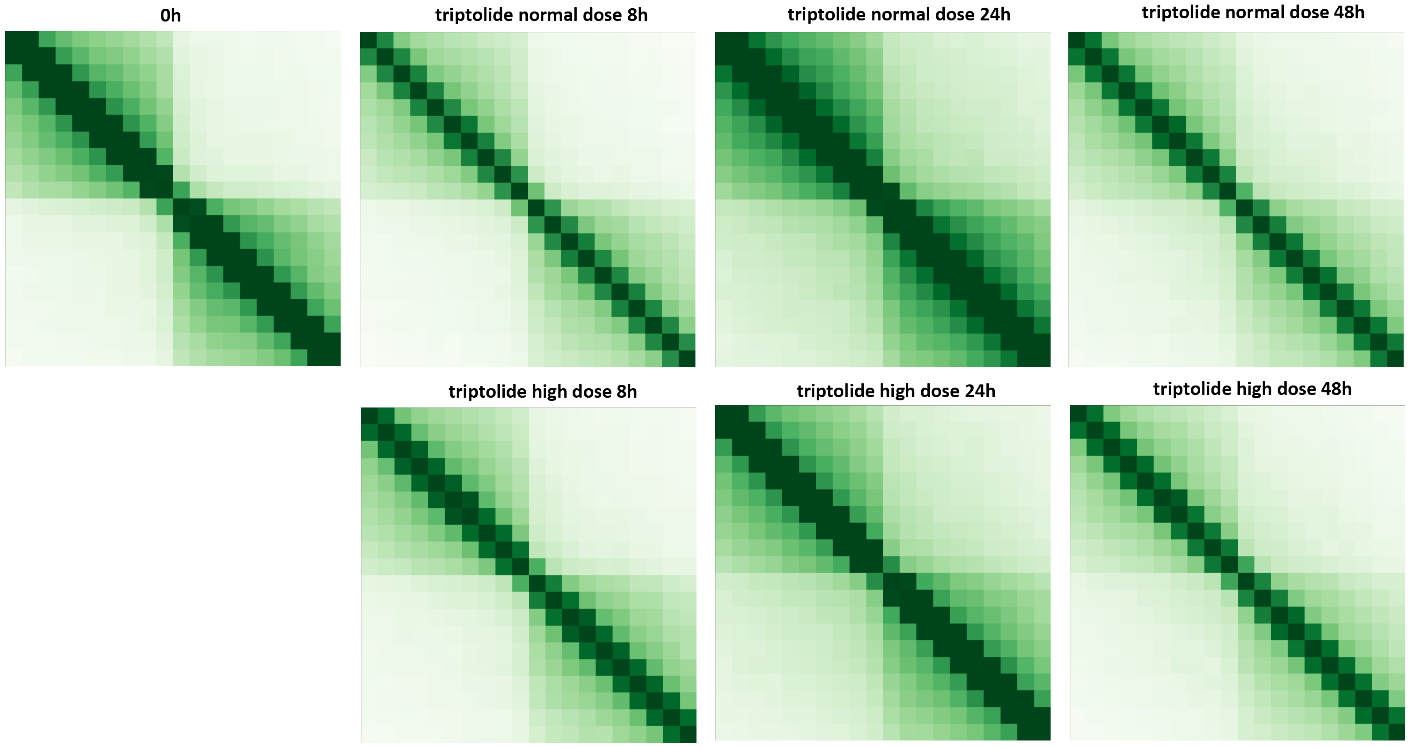


**Supplementary Figure 28: Blurring of dinoTAD boundaries upon transcriptional inhibition using triptolide**. Shown are 50-kb resolution metaplots centered on dinoTAD domain boundaries. The triptolide time course was carried out following the outline presented in Figure 2B.
